# Bro1-Mediated Trafficking Couples TOR Signalling to Cellular Metabolism and Longevity

**DOI:** 10.64898/2026.01.12.698986

**Authors:** Kristal Ng, Juhi Kumar, Aleksandra Dabrowska, Jose Clemente-Ramos, Rowshan Ara Islam, Olga Xintarakou, Anja Freiwald, Natalie Bartel, Daniela Ludwig, Despina Stamataki, John-Patrick Alao, Peter Thorpe, Markus Ralser, Michael Mulleder, Sara E. Mole, Charalampos Rallis

**Author notes:** These authors have contributed equally to this work (Kristal Ng, Juhi Kumar, Aleksandra Dabrowska, Jose Clemente-Ramos).

## Abstract

Adaptation to nutrient availability requires coordination between growth control, metabolism, and intracellular trafficking. In eukaryotes, inhibition of Target of Rapamycin (TOR) signalling robustly promotes stress resistance and longevity, yet how reduced growth signalling is coupled to organelle dynamics and proteome remodelling remains unclear. Here, we identify the conserved ESCRT-associated protein Bro1 as a central integrator of TOR signalling, vacuolar trafficking, and metabolic adaptation. Using fission yeast, we show that Bro1 is required for normal lifespan and for the global proteomic reprogramming that accompanies TOR inhibition. In Bro1 mutant cells, repression of ribosome biogenesis is uncoupled from activation of catabolic, vacuolar, and metabolic pathways, resulting in an altered metabolic state characterised by elevated lipid metabolism and increased abundance of nutrient transporters.

Mechanistically, Bro1 promotes TOR-dependent cargo deubiquitination, vacuolar trafficking, and turnover of plasma membrane hexose transporters and enables appropriate nuclear relocalisation of the transcriptional repressor Scr1. In the absence of Bro1, nutrient transporters persist at the cell surface despite TOR inhibition, conferring resistance to TOR inhibitors while impairing stress responses and reducing lifespan. Together, our findings establish Bro1 as a key coordinator linking ESCRT-mediated endosomal-vacuolar trafficking to TOR-dependent metabolic control. By coupling growth suppression to enhanced recycling and cellular maintenance, Bro1 enables the transition from growth to longevity-promoting states, revealing a mechanism connecting intracellular trafficking, metabolism, and ageing.

## Introduction

The conserved Target of Rapamycin (TOR) signalling pathway regulates stress, growth and ageing^1–3^ from yeast to human. TOR serine/threonine kinases associate with two structurally and functionally distinct multiprotein complexes termed TORC1 and TORC2^4,5^. TORC complexes coordinate basic cellular organization and metabolism^1,2,4,5^. TORC1 functions in the cytoplasm and on the lysosome^6,7^, or the lysosome-equivalent, vacuoles^8^ in fission yeast. TORC1 inhibition prolongs lifespan in all organisms tested^2,3,9–11^ through many mechanisms including the involvement of mitochondrial function^12^, autophagy^11^ and protein translation levels and fidelity^2,9,10,13^. Extension of lifespan through downregulation of protein translation depends on kinase effectors downstream of TOR^14^ (such as ribosomal S6K1) and regulation of Pol I-^15^, Pol II-^16,17^ and Pol III-dependent^14,18^ transcription. TOR inhibition and cellular growth arrest can be achieved through genetic and pharmacological treatments^14^. We have previously shown that a combination of caffeine and rapamycin inhibits the TOR pathway in fission yeast^10,19,20^. In addition, we and others have shown that the ATP-competitive TOR inhibitor torin1 effectively inhibits cell growth in fission yeast and mammalian cells^17,21^. Using a stringent screen for fission yeast mutants resistant to torin1 (20 μM) we have demonstrated that the conserved GATA transcription factor Gaf1 is fundamental for this phenotype. Beyond *gaf1* most mutants resistant to torin1 belong to the <u>E</u>ndosomal <u>S</u>orting <u>C</u>omplexes <u>R</u>equired for <u>T</u>ransport (ESCRT)^17^.

ESCRT-related proteins assemble into a multi-subunit machinery that performs topologically unique membrane bending and scission reactions. The machinery includes five distinct ESCRT complexes (ESCRT-0, -I, -II, -III and the Vps4 complex) with clear division of tasks, from interaction with ubiquitinated membrane proteins to membrane deformation and abscission^22^. The ESCRT machinery is evolutionarily highly conserved and implicated in processes required for the multivesicular body (MVB) pathway, cytokinesis and viral budding ^23^ as well as lysosome and membrane repairs^24,25^ and nuclear envelope reformation^24^. Although the pathway has expanding roles in fundamental cellular processes^22^ and has been implicated in disease^26^, knowledge on its precise connections with ageing and core metabolism in response to TOR inhibition are not fully understood^27^.

Bro1, initially implicated in the Pkc1p-MAP kinase pathway^28^, is a Class E vacuolar protein sorting (VPS) family member and an ESCRT-III adaptor protein^18^. It has subsequently been confirmed to be involved in the MVB pathway, with its association with endosomes occurring in an Snf7-dependent manner (Snf7 being the most abundant ESCRT-III core subunit)^29^ and its dissociation from the organelle requiring a functional Vps4 ATPase^30^. *Bro1* deletion mutants display accumulation of class E compartments-abnormally shaped endosomes lacking intraluminal vesicles (ILVs)-characteristic for ESCRT impairment^31^. Apart from the role in the MVB pathway, Bro1 has an indirect function in deubiquitination of cargos through recruitment of Doa4, a ubiquitin hydrolase^32^, to endosomes. More recently, Regulator for free Ubiquitin Chains 1 (Rfu1) has also been shown to be recruited by Bro1, thus, confirming further its role in ubiquitin homeostasis^33^. As Doa4 also functions as a negative ESCRT-III membrane scission activity regulator, binding of Bro1 to Vps20 (Doa4 inhibitor) further promotes Doa4 function by allowing its association with Snf7, regulating budding of the ILVs^34^. Its role in late endosome formation is also dependent on its ability to stimulate Vps4 ATPase activity which is required for membrane budding^35,36^.

Here, we show that Bro1 is required for vacuolar fusion events, normal chronological lifespan and for lifespan extension mediated by TOR inhibition. Using gene-gene-drug genome-wide assays we reveal genetic combinations that sensitize *bro1* mutant cells. Absence of Bro1 results in high levels of glucose and other transporters. Torin1-mediated Ght5 hexose transporter internalization and recycling still occur in *bro1* mutants, but not efficiently. *Tor1* deletion sensitizes *bro1* mutants to TOR inhibition. Bro1-dependent high Ght5 expression levels occur due to the cytoplasmic retainment of the transcriptional repressor Scr1. Our results reveal circuits that modulate metabolism and lifespan and are relevant to cancer biology as they provide mechanistic insights into how cells can escape TOR inhibition and promote growth programs.

## Results

### *Bro1* lifespan effects and TOR activity-dependent genetic interactomes

We have previously shown that cells mutant for *gaf1* are resistant to torin1 and short-lived. We, therefore, examined the chronological lifespan (CLS) of *bro1*Δ cells in EMM2 media as previously described ^17^. Indeed, mutant cells are short-lived with median and maximum lifespans of 2 and 7 days, respectively, as opposed to 4 and 8 days of *wt* cells (log rank, p=2.9×10^-4^) (Fig. 1A). We wondered whether *bro1* is required for TOR-inhibition-mediated lifespan extension. We repeated our CLS assay following treatment of *wt* and *bro1*Δ cells with 2µM of torin1 at OD_600_ 0.5. This concentration of the drug was chosen to delay but allow the growth of the yeast cells to stationary phase rather than block their cell cycle progression completely. While *wt* cells showcase increased lifespan (log rank p=2.5×10^-4^) with median and maximum values of 5 and 10 days, respectively, *bro1*Δ cells do not exhibit statistically significant change (n.s., log rank, p=0.25) (Fig. 1A). AUC analyses of obtained lifespan curves further confirm these results (Fig. 1B). Taken together, our results support that *bro1* is required for both normal lifespan and for torin1-mediated lifespan extension.

**Figure 1.**
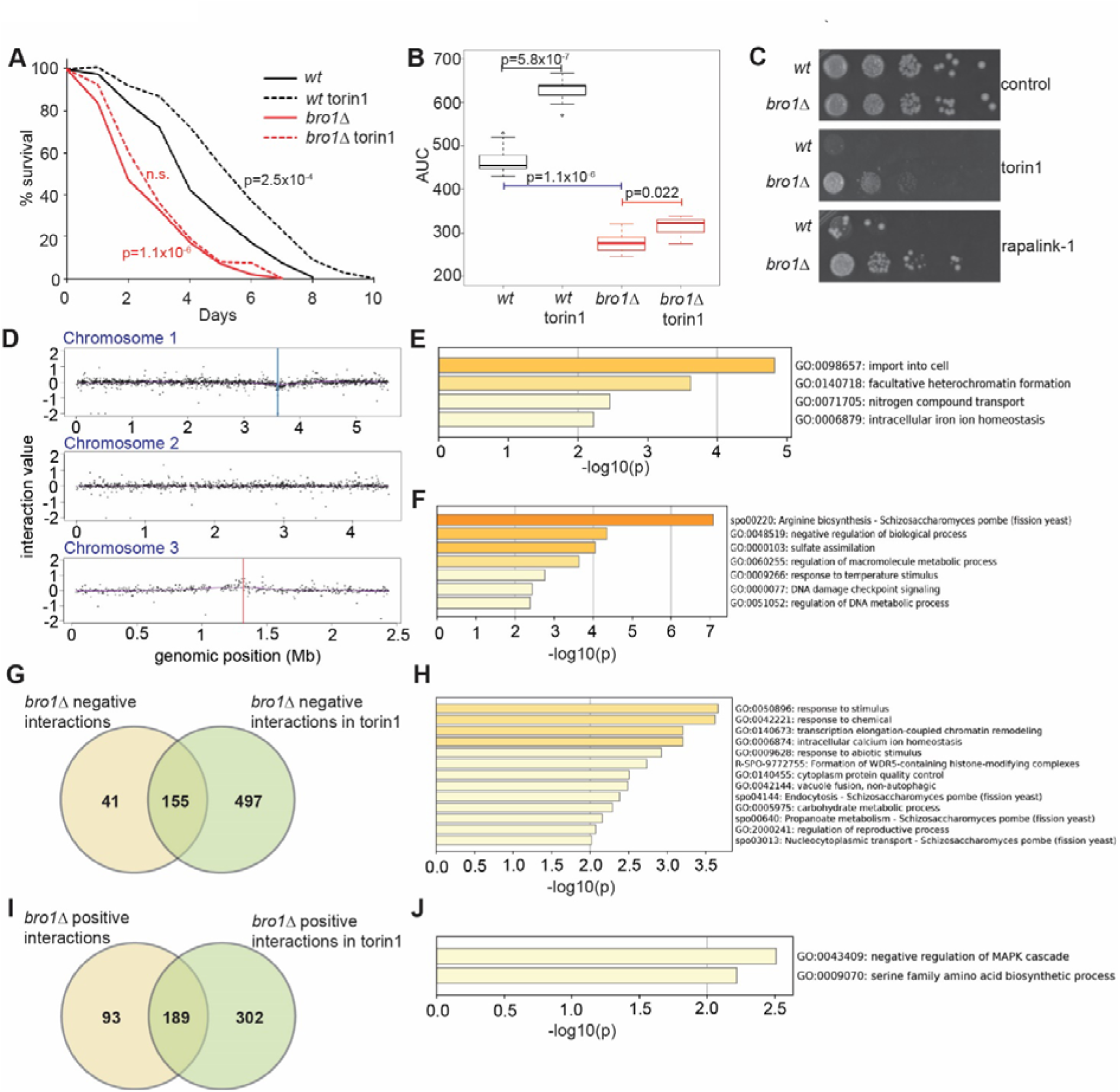
*Bro1* genetic interactomes and chronological lifespan in mTOR-active and inactive states. **A.** Chronological lifespan assays for *wt* and *bro1*Δ cells in EMM2 media in the presence and absence of torin1 as previously described ^17^. Assays have been performed in 3 biological replicates with 3 technical replicates each (n=9 for each timepoint assessed). P-values indicate log rank tests. n.s.: not significant difference between *bro1*Δ and *bro1*Δ with torin1. Black font p-value corresponds to statistics between *wt* and *wt* with torin1 while red font p-value corresponds to statistics between *wt* and *bro1*Δ cells. **B.** Area under the curve (AUC) measurements for each genotype and condition in **A** (n=9). Students t-test has been performed between indicated pairs with p-values shown in the panel. **C.** Serial 10-fold dilutions of fast growing OD_600_ 0.5 *wt* and *bro1*Δ cells on agar plates containing compounds that inhibit TOR as indicated (5µM torin1 and 100nM rapalink-1). **D.** Schematic showing mapping of global genetic interactions of *bro1* in the absence and absence of torin1. **E.** GO enrichment analyses for positive genetic interactions of *bro1.* **F.** GO enrichment analyses for negative genetic interactions of *bro1* **G.** Venn diagram showcasing *bro1* negative interactions that only happen in the presence of torin1. **H.** GO enrichment analysis for negative *bro1* genetic interaction gene lists in the presence of torin1. **I.** Venn diagram showcasing *bro1* positive interactions that only happen in the presence of torin1. **J.** GO enrichment analysis for positive *bro1* genetic interaction gene lists in the presence of torin1.

*bro1* mutants are resistant to the ATP-competitive TOR inhibitor torin1 ^17^. To further validate the resistance result and to determine whether *bro1* mutant cells are resistant to a different pharmacological TOR inhibition we have performed standard spot assays in the presence of torin1^17^ (ATP-competitive imhibitor) and rapalink-1^37^ (that combines rapamycin with an ATP-competitive inhibitor). Wild-type and mutant cells were grown to OD_600_ 0.5 and then serially diluted 10-fold and spotted on YES agar media and within the required conditions. While the growth of *wt* cells is greatly inhibited, *bro1*Δ cells in mTOR inhibited conditions divide in a comparable manner to untreated cells (Fig. 1C). To further our understanding of the roles of *bro1* and investigate deeper the genetic basis of *bro1*Δ mutant cells in TOR signalling resistance we performed Synthetic Genetic Arrays (SGAs) in the absence and presence of torin1. SGA is a cell-based genome-wide approach that can identify genetic interactions for a gene of interest with ∼3,400 mutants (corresponding to non-essential fission yeast genes) ^38^ through the systematic construction of double mutants. Estimating the fitness of single mutants and their corresponding double mutants enables the quantitation of genetic interactions, distinguishing negative (i.e. synthetic sick or lethal) and positive (within pathway and suppression) interactions. We have isolated 196 negative interactions and 282 positive interactions throughout the genome (Fig. 1D, Table S1) with z-score cutoffs -0.15 and 0.15 respectively^19^. Negative interactions are enriched^39,40^ (Gene Ontology Biological Process) in genes encoding for plasma membrane transporters (GO: import into cell) including amino acid transporters (GO: amino acid compound transport), heterochromatin formation and intracellular iron homeostasis (Fig. 1E). Positive interactions are enriched in the KEGG pathway arginine biosynthesis and in Biological Process GO terms regulation of metabolic process (including several ribosomal subunits), DNA metabolic process, transcriptional regulators including Pcr1 and Zip1, the co-repressor Tup1, Clr6 histone deacetylase components as well as several ubiquitin ligases and factors involved in vesicular transport (COP and ESCRT complexes).

To uncover gene combinations that sensitise *bro1*Δ mutant cells to torin1 we repeated the SGAs in the presence of 5 µM of the drug (Table S2). As *bro1*Δ cells are torin1 resistant they render most of the library resistant too (Fig. S1). Firstly, using the same z-score cutoffs as in the SGAs (−0.15 and 0.15 for negative and positive interactions, respectively) we have isolated 652 negative *bro1* genetic interactions in the presence of torin1 that are enriched in vesicle-mediated transport, vacuole fusion and chromatin remodelling among others (Fig. S2A). A large portion of these genes (97/652) are core environmental stress response genes^41^ (induced in all stresses). The 491 positive interactions are enriched in glutamine biosynthetic process, proteasome assembly, negative regulation of translation and ribosome genes (Fig. S2B). To identify mutants that sensitise *bro1*Δ cells in torin1 we compared negative interactions in the absence and presence of torin1 towards distilling the mutants that have such interactions (or even become synthetic lethal) with *bro1* only in the presence of the drug. 497 mutants show negative interactions with *bro1* in torin1 (Fig. 1G). These are enriched in several GO terms, notably chromatin remodelling, intracellular calcium homeostasis, protein quality control, non-autophagic vacuole fusion and endocytosis (Fig. 1H). The last 2 categories have the largest torin1 sensitising effect to *bro1* mutant background and include ubiquitin ligases, amino acid and other nutrient transporters and *isp7* that encodes for a regulator of amino acid transporters^42^.

Transporters and phagosome-related protein categories are also predicted to be physical interactors of Bro1 protein using our machine learning tool Pint^43^ (Fig. S2C, Table S3). We then wondered what would make *bro1*Δ cells even more resistant to TOR inhibition. Comparing SGA positive interactions in the absence and presence of the drug revealed 302 mutants that do this (Fig. 1I). Enrichment analysis uncovers negative regulation of MAPK cascade and serine amino acid family biosynthetic process (Fig. 1J). Further enrichment analysis using the fission yeast-specific Angeli tool^39^ shows that 33 out of the 302 genes are involved in cytoplasmic translation and include *ssp2* (coding for the AMPK catalytic subunit), *hri2* (coding for the eIF2 alpha kinase Hri2) and several ribosomal proteins. Many of these genes are reported to be required for survival in stationary phase^44^ as we do, here, for *bro1* (Fig. 1A). Our gene-gene and gene-gene-drug analyses reveal the *bro1* interactome map and point to genes related to nutrient transporters, endocytosis and vacuolar fusion as the ones that can sensitise *bro1*Δ cells to torin1.

### *Bro1* is required for vacuole fusion events and related amino acid homeostasis

Although not part of the ESCRT complexes, Bro1 is implicated in cargo processing for vacuoles and endosomes. Given our results following SGAs, we wondered whether absence of Bro1 affects fission yeast vacuolar dynamics and related functions such as amino acid concentration regulation^45^. Towards this, we have stained *wt* and *bro1*Δ cells with 7-amino-4-chloromethylcoumarin (CMAC) without and with torin1 treatment (5 µM for 2 hrs) as well as following osmotic shock through treatment with water (2 hrs). CMAC contains a mildly reactive chloromethyl moiety that reacts with accessible thiols on peptides and proteins to form an aldehyde-fixable conjugate and accumulates in the vacuole lumen^46^. In control states we observed that *bro1*Δ vacuoles may be smaller or fragmented compared to those of normal cells (Fig. 2B and 2D). Such vacuoles were no longer visible following treatment with torin1 (Fig. 2C and 2E). Quantifications of these results showed that *bro1*Δ vacuoles are significantly smaller in diameter and area while torin1 treatment restores the phenotype to *wt* appearance (Fig. 2F and 2G). Notably, the differences between *wt* and *bro1*Δ cells in control states are probably underestimated: we tried several scripts and approaches that failed to provide measurements for the very small CMAC-positive formations representing a fragmented vacuole or endosome moiety. To test vacuolar dynamics in the absence of *bro1* we subjected cells to osmotic shock by transferring them in water for 2 hrs. In these conditions, vacuoles fuse as seen in *wt* cells (Fig. 2H and 2I) with the presence of large CMAC-positive vacuolar formations, quite evident. Contrary to *wt*, *bro1*Δ cells fail to undergo vacuole fusion events (Fig. 2J and 2K). Quantifications of these effects clearly demonstrate the increase in both diameter and area of *wt* vacuoles in water with the phenomenon completely suppressed within *bro1*Δ cells. Taken together, our data show that *bro1* is required for normal vacuolar size and remodelling.

**Figure 2.**
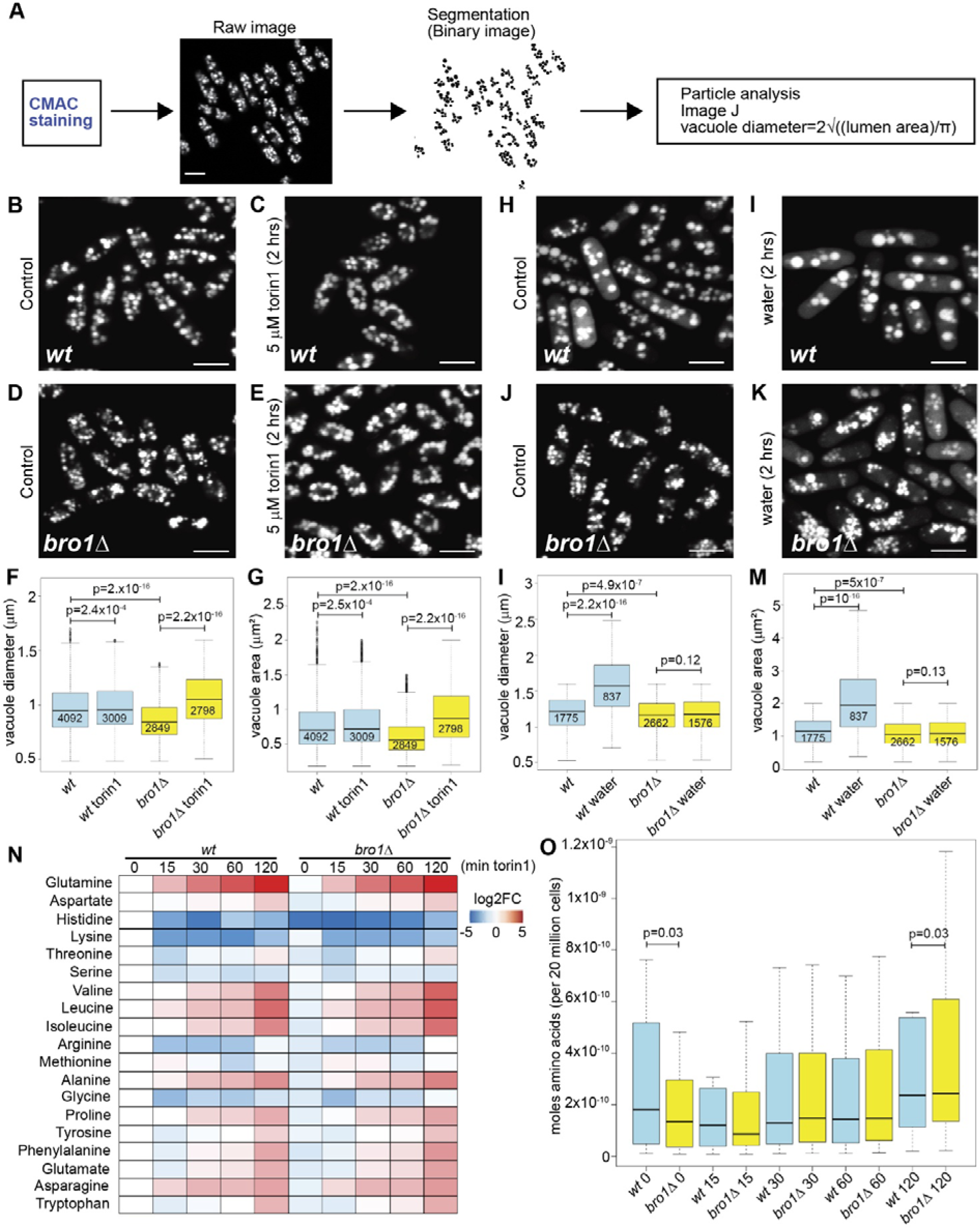
*Bro1* is required for vacuolar dynamics and intracellular amino acid levels. **A.** Schematic of vacuole size analysis flow utilised. Cells were grown and treated in YES media. **B-E.** Representative images of CMAC stained *wt* (**B**, **C**) and *bro1*Δ (**D**, **E**) cells in control (DMSO-only) or 5 µM torin1-treatment (for 2 hrs) conditions, as indicated. **H-K.** Representative images of CMAC stained *wt* (**H**, **I**) and *bro1*Δ (**J**, **K**) cells in control (YES) or water-treated (for 2 hrs) conditions as indicated. Bar in **B**-**K**: 10 µm. **F.** Vacuole diameter quantifications in conditions and genetic backgrounds as indicated. **G.** Vacuole area quantifications in the same conditions as in F. **L.** Vacuole diameter quantifications for *wt* and *bro1*Δ cells in control and water treatments as indicated. The differences in measurements between control states in panels F and L are attributed to DMSO. **M.** Vacuole areas in conditions as in L. The differences in measurements between control states in panels G and M are attributed to DMSO. For all boxplots numbers within boxes are the n numbers of quantified vacuoles in each case. P-values indicate Wilcoxon statistical testing for the vacuole populations, as indicated. Data for all quantifications derives from three independent biological replicates. **N.** Heatmap for the quantification of intracellular amino acid concentrations in *wt* and *bro1*Δ cells through HILIC-MS/MS in a time course treatment with torin1 (0-120 min). All values are normalised to *wt* at time point 0 min. **O.** Boxplots indicating the intracellular amino acid contents distribution across all conditions shown in N. P-values (0.03 in both cases as shown) indicate paired students t.test statistical testing.

Vacuoles are highly regulated structures and serve as main storage compartments for basic and neutral amino acids^47^. We have shown that *bro1*Δ cells showcase smaller vacuoles that are restored to size ranges comparable to those of *wt* cells following torin1 treatment. We, therefore, wondered whether the observed differences in structure and size reflect amino acid content and whether the morphological ‘rescue’ through torin1 is also accompanied by amino acid content restoration. We have performed intracellular amino acid profiling through mass spectrometry (HILIC-MS/MS) as previously described^48^. Indeed, *bro1*Δ cells contain less intracellular amino acids, nevertheless, torin1 treatment increases amino acid content in both genetic backgrounds and, importantly, restores amino acid content in mutant cells (Figure 2N).

Quantifications demonstrate a total amino acid content of 1.15X10^-8^ mol per 20 million *wt* cells as opposed to 8.49×10^-9^ mol in *bro1*Δ cells (paired student’s t.test p=0.03) with median amino acid content values of 1.82×10^-10^ and 1.34×10^-10^ mol respectively. Following torin1 treatment the content increase to 4.24×10^-8^ and 4.98×10^-8^ mol per 20 million cells (paired student’s t.test p=0.03) with median values at 2.37×10^-10^ and 2.43×10^-10^ mol, respectively (Fig. 2O). Our data show that fast-growing *bro1*-mutant cells store less amino acids compared to *wt* cells but are restored following mTOR inhibition.

### Absence of *bro1* affects gene expression in normal and mTOR inhibition states

As Bro1 affects both lifespan, resistance to torin1 and rapalink1 and vacuolar appearance and dynamics we wondered whether its absence affects gene expression in both normal and TOR-inhibition states. Towards this fast-growing cells were treated with 5 µM torin1 or a combination of 100 µM rapamycin and 10mM caffeine^10^. *Bro1* mutants show minimal differential gene expression compared to *wt* cells (Fig.3A) with 10 genes upregulated and 11 downregulated in a 2-fold cutoff (Table S4 for differentially expressed gene and Table S5 for all RNAseq data). One of the genes among the upregulated is *ght5*, a hexose transporter. Examination of gene expression correlations in both untreated and torin1 and caffeine-rapamycin treated cells (for 2 hrs and 5 hrs) reveal that *bro1*Δ mutants have similar responses to the drugs as normal cells (Fig. 3B), while Principal Component Analysis (PCA) analysis can distinguish between the treatment regimes with PCA1 accounting for 61.11% of variability and PCA2 9.74%. Torin1 imposes considerable changes in gene expression in both normal and *bro1*Δ cells (Fig.S3 and Fig.S4, respectively). GO ontology enrichment analyses show that catabolic processes, transmemberane transport among other categories are upregulated in both cases while anabolic processes such ribosome biogenesis and protein translation are downregulated, as expected (see Fig.S3 and Fig.S4 for corresponding volcano plots and GO enrichments) and in agreement with previous results from our group based on expression microarrays^17^. Caffeine-rapamycin pairwise comparisons at 2-fold cutoffs reveal small number of differentially expressed genes in the 2 hrs and 5 hrs timeframes (Fig.S5). Although upregulated genes are related to catabolism and downregulated genes to anabolism (TORC1 related processes), GO enrichments were not satisfactory in the number of categories present in these limited gene lists. We therefore performed further comparisons with 1.5-fold cutoffs for both *wt* (Fig.S6) and *bro1*Δ cells (Fig.S7). GO in upregulated genes in the 2 hrs timepoint for *wt* cells are related to stress response and mitophagy while downregulated genes are related to ribosomal biogenesis and processing, amino acid catabolic process, and iron import into cell (Fig.S6). The latter category is also present at the later timepoint (5 hrs). Catabolic processes and responses to stresses categories are also upregulated in the 2 hrs time point for *bro1*Δ while ribosome biogenesis and translation as well as iron import is present in the downregulated categories (Fig.S7). Nevertheless, following 5 hrs treatment with caffeine-rapamycin cells mutant for *bro1* show more changes in gene expression compared to the *wt* (see Fig.3C and Fig.S7 D-I and comparison with Fig.S6 D-I).

**Figure 3.**
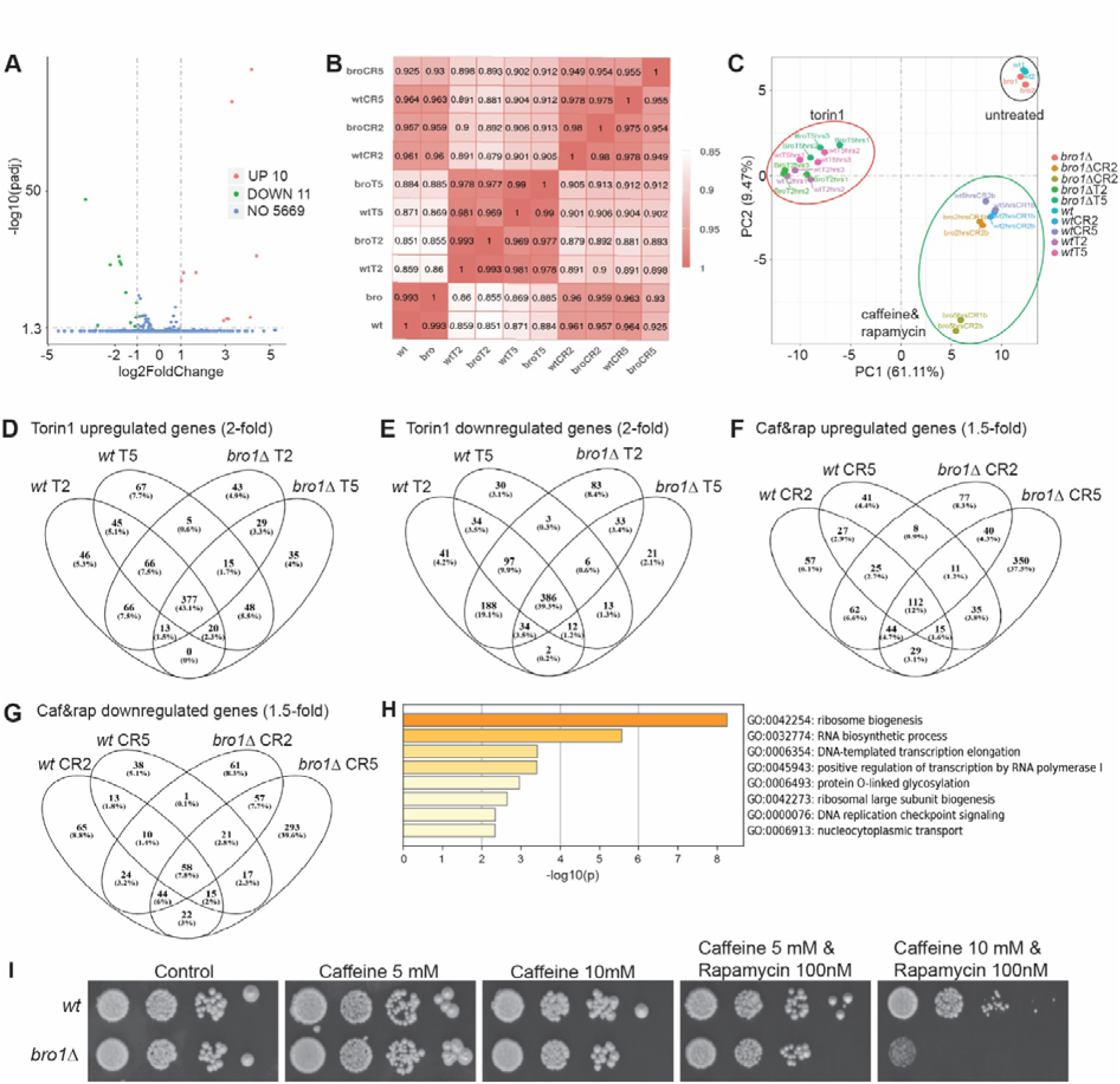
*Bro1* affects gene expression in mTOR-active and mTOR-inhibition states. **A.** Volcano plot for differentially expressed genes between *wt* and *bro1*D cells. **B.** Correlation matrix between RNA-seq samples from normal (*wt*) and *bro1*D (bro) cells without treatment as well as treated with torin1 (T) or caffeine-rapamycin for 2 or 5 hrs as indicated. **C.** Principal Component Analysis (PCA) for the samples analysed through RNAseq. **D.** Venn diagram showing overlaps for 2-fold upregulated genes in *wt* and *bro1*D mutants treated with torin1 (T) for 2 or 5 hrs as indicated. **E.** Venn diagram showing overlaps for 2-fold downregulated genes in *wt* and *bro1*D mutants treated with torin1 (T) for 2 or 5 hrs as indicated. **F.** Venn diagram showing overlaps for 1.5-fold upregulated genes in *wt* and *bro1*D mutants treated with caffeine-rapamycin (CR) for 2 or 5 hrs as indicated. **G.** Venn diagram showing overlaps for 1.5-fold downregulated genes in *wt* and *bro1*D mutants treated with caffeine-rapamycin (CR) for 2 or 5 hrs as indicated. **H.** Serial 10-fold dilutions of fast growing OD_600_ 0.5 *wt* and *bro1*Δ cells on agar plates containing caffeine and rapamycin in combinations and concentrations as indicated.

To directly understand the differences between the responses to the drugs between *wt* and *bro1*Δ we compared differentially expressed genes followed by GO enrichments in the gene lists unique for the two genotypes. 158 genes are upregulated exclusively in the *wt* following torin1 treatment (for both 2 and 5 hrs, Fig.3D). These are enriched for amide transport and carbohydrate catabolic process. 107 genes are exclusively upregulated in *bro1*Δ cells (for both 2 and 5 hrs, Fig.3D). No GO enrichments are observed in this gene list, nevertheless, several ncRNAs and membrane transporters are included. 105 genes are downregulated in *wt* cells following torin1 treatment (for both 2 and 5 hrs, Fig. 3E). These are enriched for protein import into mitochondrial matrix and transmembrane transport. 137 genes are downregulated in *bro1*Δ cells, enriched for metabolism of lipids and 5.8S rRNA maturation. Taken together, our results show that while transmembrane transport genes are downregulated in normal cells, in the absence of *bro1* these genes are unaffected while many other membrane transporters are instead, upregulated.

We then examined differentially expressed genes in response to caffeine-rapamycin that contrary to torin1 that affects both TORC1 and TORC2, targets primarily TORC1 in fission yeast^10^. 125 genes are exclusively upregulated in *wt* cells (1.5-fold, both 2 and 5 hrs timepoints, Fig.3F) with 12 related to vesicle-mediated and 6 to transmembrane transport. Notably genes encoding for ESCRTIII complex are present in this category. 467 genes are upregulated in in *bro1*Δ cells (1.5-fold, both 2 and 5 hrs timepoints) with 350 of them in the 5 hrs timepoint. These are enriched in vesicular transport (related to ER and Golgi), membrane organisation, nucleophagy and autophagy among others (Fig.3F). We then examined the downregulated genes following caffeine-rapamycin treatment. 116 genes are downregulated exclusively in *wt* cells (1.5-fold, both 2 and 5 hrs timepoints, Fig.3G) and examination of GO enrichment indicates postmitotic nuclear pore complex reformation and alcohol metabolic process as enriched categories. 12 of these genes are related to transcription 8 to chromatin organisation and 5 to transmembrane transport. On the other hand, 411 genes are downregulated in *bro1* mutant cells (1.5-fold, both 2 and 5 hrs timepoints) with 294 only for the 5 hr timepoint (Fig.3G). Analysis of these gene lists demonstrates a large enrichment on ribosome biogenesis, Pol I-related transcription and RNA biosynthetic process also related to ribosomal RNA and ribosomal proteins (Fig.3H). Our data indicate that caffeine-rapamycin treatment has a notable effect on *bro1*Δ compared to *wt* cells and might signify that cells lacking *bro1* while resistant to torin1 (Fig.1) might be more sensitive to caffeine-rapamycin treatment. To test this hypothesis, fast-growing (OD_600_=0.5) *wt* and *bro1*Δ were serially diluted and spotted on YES plates containing 100nM rapamycin, 5mM and 10 mM caffeine and a combination of the drugs in these concentrations. Indeed, *bro1*Δ mutant cells are sensitive to caffeine-rapamycin in 10mM caffeine and 100nM rapamycin, concentrations previously tested and used for genome-wide screens^10,19,20^.

### Bro1 couples TOR inhibition to proteomic and metabolic reprogramming

Given the defects in TOR signalling, vacuolar function, and stress resilience observed in *bro1*Δ cells, we next examined how loss of Bro1 impacts on global proteome remodelling in response to TOR inhibition. Quantitative proteomic profiling of fast growing (OD_600_=0.5) wild-type and *bro1*Δ cells under basal (untreated) conditions and following torin1 treatment, revealed strong separation by genotype and treatment in principal component analysis (Figure 4A). PCA1, accounting for 60.7% of the variance, primarily distinguished *bro1*Δ from wild-type cells, consistent with the constitutive transcriptional and signalling alterations described above. PCA2 (21.8% variance) reflected torin1 exposure, indicating that Bro1 loss substantially reshapes the proteomic response to TOR inhibition (summary of proteomics results is presented in Table S6). Interestingly, the difference between *wt* and *bro1* mutant cells is reduced upon treatment, a result reminiscent of the vacuolar and amino acid content rescues observed in Fig.2.

**Figure 4.**
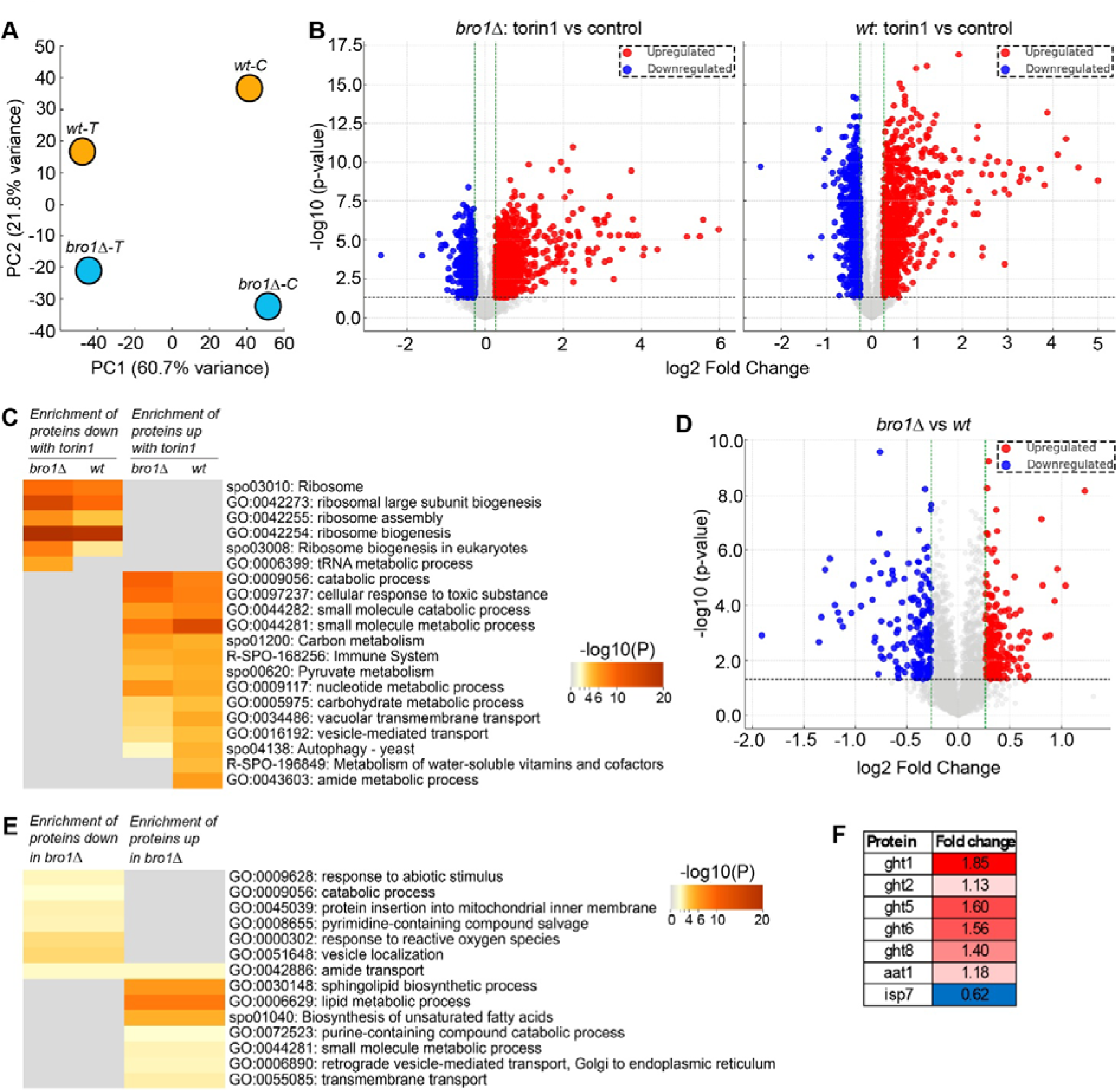
Bro1-dependent proteomic and metabolic reprogramming upon TOR inhibition. **A.** Principal component analysis (PCA) of quantitative proteomics data from wild-type (*wt*) and *bro1*Δ cells grown in control conditions (C) or treated with torin1 (T). PCA1 and PCA2 explain 60.7% and 21.8% of the variance, respectively, demonstrating clear separation by genotype and TOR inhibition status. **B.** Volcano plots showing patterns of differentially expressed proteins in torin1 vs intreated conditions for normal and *bro1* mutant cells. **C.** Gene Ontology (GO) and pathway enrichment analysis of proteins significantly downregulated (left) or upregulated (right) following torin1 treatment in *wt* and *bro1*Δ cells. Enrichment scores are displayed as -log10(P). Downregulated categories are dominated by ribosome biogenesis, ribosomal assembly, and tRNA metabolic processes, whereas upregulated categories include catabolic processes, central carbon metabolism, vacuolar transport, autophagy, and stress-response pathways. **D.** Volcano plot for differentially expressed proteins in *bro1* mutant cells vs normal cells. **E.** GO enrichment analysis of proteins significantly decreased (left) or increased (right) in *bro1*Δ cells relative to *wt* cells under control conditions. *bro1*Δ cells show reduced representation of stress response, vesicle localization, and mitochondrial-associated processes, alongside increased enrichment of lipid, sphingolipid, and small-molecule metabolic pathways. **F.** Fold-change values for selected plasma membrane transporters and metabolism-related proteins differentially abundant in *bro1*Δ cells relative to *wt*. Increased abundance of multiple hexose transporters (Ght1, Ght2, Ght5, Ght6, Ght8) is observed, while the amino acid transporter regulator Isp7 is reduced. Values represent mean fold changes derived from quantitative proteomics.

In wild-type cells, torin1 treatment triggered a canonical TOR inhibition signature characterised by coordinated downregulation of ribosome biogenesis, ribosomal assembly, and tRNA metabolic pathways, alongside induction of catabolic, metabolic, and vacuolar processes (Figure 4B, C). These changes parallel the autophagy induction and vacuolar remodelling observed following TOR inhibition in earlier experiments (Figures 2 and 3). In *bro1*Δ cells, however, this adaptive response was altered. Although ribosome-associated pathways were still suppressed upon torin1 treatment, enrichment of metabolic and vacuolar pathways was redistributed, consistent with impaired execution of TOR-dependent proteome reprogramming in the absence of Bro1 (Fig.4B, C).

Analysis of basal proteomic differences further revealed that *bro1*Δ cells exhibit a distinct metabolic state even under nutrient-replete conditions. Proteins involved in stress responses, vesicle localisation, mitochondrial inner membrane protein insertion, and reactive oxygen species handling were reduced (172 proteins, <1.2-fold, p-value<0.05, Figure 4D, E), whereas lipid metabolism, sphingolipid biosynthesis, unsaturated fatty acid synthesis, and small-molecule metabolic processes were enriched (179 proteins >1.2-fold, p-value<0.05, Figure 4D, E). This constitutive metabolic rewiring is consistent with the altered TOR activity, vacuolar trafficking defects, and reduced lifespan previously observed in bro1Δ cells (Figures 1 and 2). Consistent with these global changes, *bro1*Δ cells displayed increased abundance of multiple plasma membrane hexose transporters, including Ght1, Ght2, Ght5, Ght6, and Ght8, together with reduced levels of the nutrient-responsive regulator Isp7^42^ (Figure 4F). This transporter profile suggests enhanced nutrient uptake and dysregulated nutrient sensing, providing a mechanistic link between Bro1-dependent vacuolar trafficking, aberrant TOR signalling, and the failure to appropriately transition from anabolic growth to catabolic maintenance states.

Collectively, these data position Bro1 as a critical regulator of TOR-dependent proteomic and metabolic adaptation, coordinating vacuolar function and protein recycling with nutrient sensing and cellular homeostasis. Impaired proteome remodelling in *bro1*Δ cells likely underlies their defective stress responses and reduced longevity described earlier in the manuscript.

### *Bro1* regulates homeostasis and TOR inhibition resistance through transporter expression and recycling

Many genes upregulated in the *bro1* mutant are clustered together within the genome (on chromosome I, Fig.S8). Promoters of these genes together with the gene encoding the main hexose transporter in fission yeast, Ght5 (located in chromosome II), are shown be bound by the transcriptional repressor Scr1^49^, a protein also predicted through our AI tools to be physically interacting with Bro1^43^ (Fig.S2C, Table S3). We therefore examined whether Scr1 localisation is affected in the *bro1* mutant cells in both normal and torin1-treatment conditions using a strain in which Scr1 is tagged with Green Fluorescent Protein (GFP, *Scr1-GFP*). In *wt* cells Scr1 is found in both the nucleus and the cytoplasm (45% in nucleus and 55% in cytoplasm, Fig. 5A). Nevertheless, in *bro1*Δ cells Scr1 is located outside of the nucleus on membrane-bound formations, most possibly vacuoles. Torin1 treatment results in 100% nuclear localization of Scr1 in *wt* cells from 15 min (Fig. 5A) while nuclear translocation in *bro1*Δ cells is impaired with 0% localization in 15 min and 42% after 2 hrs, the latter resembling the conditions within untreated *wt* cells. Given that Scr1 is predicted to interact with Bro1 (Fig. S8B) and that global ubiquitination is elevated in *bro1*Δ mutant cells (Fig.5B), we wondered whether Bro1 affects ubiquination levels of Scr1 apart from its subcellular localization. Towards this, we performed immunoprecipations using α-GFP antibody and extracts from *Scr1-GFP* and *Scr1-GFP bro1*Δ strains. While we have used the same total protein amounts for immunoprecipitations (40 µg total protein) we detect reduced amounts of Scr1-GFP protein in the absence of *bro1.* Although Scr1 levels are lower, its ubiquitination is evidently higher (Fig. 5C). Our results indicate that Bro1 affects the protein levels as well as the ubiquitination of Scr1. This can explain the upregulation of *ght5* transcripts in *bro1*Δ cells (Fig.3A, Table S1) and the increased abundance of Ght proteins in our proteomic data (Fig.4D). Beyond the expression levels of Ght5 we then wondered whether Bro1 is important for Ght5 localisation within the cell. Using the Ght5-GFP strain, we observe that in normal conditions Ght5 is located mainly on the cell membrane. Torin1 treatment results in the internalisation of Ght5 within 5 min and in its localisation to vacuoles or vesicles within 15 min (Fig.5D). Nevertheless, in the *bro1*Δ background, Ght5-GFP is more abundant on the membrane (through qualitative observation) in agreement with our western blot results. Following torin1 treatment, we do observe internalisation and vacuole-associated Ght5. Nevertheless, its presence on the plasma membrane persists (Fig.5D). This may explain the reason that *bro1*Δ cells are resistant to torin1: upon TOR inhibition, while glucose transportation may be reduced, this is only repressed at a level where cells have more than adequate glucose levels to support their growth.

**Figure 5.**
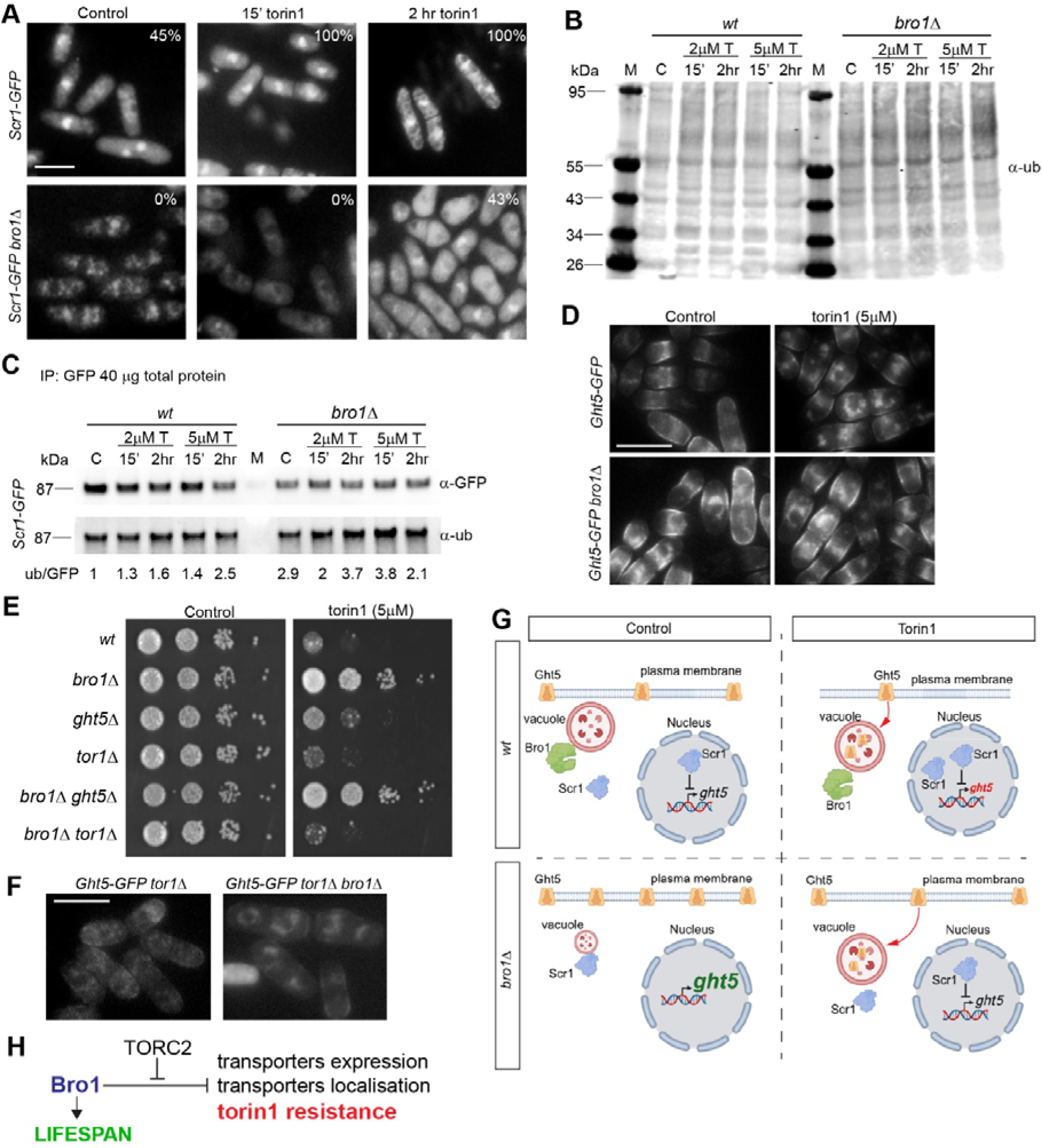
Bro1 links TOR inhibition to transporter ubiquitylation, vacuolar trafficking, and nutrient-dependent survival. **A.** Scr1-GFP localization using live-cell microscopy on normal (*Scr1-GFP*) and *bro1* mutant cells (*Scr1-GFP bro1*D) in control and torin1 treatments for 15 min (15’) and 2 hours (2 hr) as indicated. The percentage of cells showing Scr1 localization into the nucleus for every condition examined is provided in the top right corner of each panel. Scale bar 10 µm. **B.** Western blot against ubiquitin using 40 µg protein extracted from *wt* and *bro1*D cells in control (C) and torin1 (T) 2 µM and 5µM treatments for 15 min (15’) and 2 hours (2 hr). M: protein marker. **C.** Western blot analysis following immunoprecipitation of Scr1-GFP with α-GFP antibody from 40 µg total protein extracted from normal (*Scr1-GFP* strain labelled as *wt*) and *bro1*D (*Scr1-GFP bro1*D strain labelled as *bro1*D). IP samples are blotted for GFP (α-GFP) and ubiquitin (α-ub). Numbers under ubiquitin blot shows the ratios of ub/GFP normalized for the control of *wt.* **D.** Fluorescence microscopy of Ght5–GFP in wild-type and *bro1*Δ cells under control conditions or following torin1 treatment (5 µM). In wild-type cells, TOR inhibition promotes vacuolar trafficking of Ght5, whereas *bro1*Δ cells retain Ght5 at the plasma membrane. Scale bar, 10 µm. **E.** Spot dilution growth assays of the indicated genotypes on control medium or medium containing torin1 (5 µM). Loss of Bro1 confers resistance to TOR inhibition, which is suppressed by deletion of *tor1*. **F.** Fluorescence microscopy of Ght5–GFP in *tor1*Δ and *tor1*Δ *bro1*Δ cells. Ght5 is found in vacuoles in *tor1* mutants as previously reported^79^ while cytoplasmic localisation is maintained in *tor1*Δ *bro1*Δ. Scale bar, 10 µm. **G.** Model for Bro1 role on vacuole size, Scr1 localization and Ght5 expression that affects metabolism, lifespan and resistance to TOR inhibition. Red-font Ght5 signifies low expression while large green-font Ght5 signifies high expression levels. **H.** Summary model linking Bro1 to TORC2-dependent regulation of transporter expression, transporter localisation, torin1 resistance, and lifespan.

Our SGA analysis has revealed that *bro1* genetically interacts with genes coding for endocytosis proteins and transporters. In addition, interactomes of *bro1* in the presence of torin1 showed that plasma membrane transporters deletions can sensitise *bro1*Δ cells (Fig.1). We therefore examined whether *ght5* deletion can sensitise *bro1* mutant cells in torin1. Fast growing (OD_600_=0.5) *wt*, *bro1*Δ, *ght5*Δ and *bro1*Δ *ght5*Δ cells were serially diluted and spotted on plates containing 5 µM torin1. Nevertheless, *ght5* deletion does not significantly sensitise *bro1* mutant cells in this spot assay (Fig.4E). This is probably due to the upregulation of other Ght transporters (Fig.4D). The difference with the SGA approach is that the latter provides quantitative results while even small fitness differences can be detected. As TORC2 is required for transporters’ localization and internalization we, then, wondered whether *tor1* (core component of fission yeast TORC2) deletion can affect *bro1* mutant cells’ resistance to torin1. Indeed, *tor1* mutant cells are sensitive to torin1 treatment and render *bro1*Δ sensitive to the drug. Moreover, Ght5-GFP is mislocalised in *tor1*Δ cells and found within the cytoplasm (possibly in endosomes) and not on the plasma membrane (Fig.5F). In the absence of both *tor1* and *bro1* Ght5-GFP while enhanced compared to *tor1* single-mutant cells, is still found in the cytoplasm (perinuclearly, most possibly within the endoplasmic reticulum) (Fig.5E). In summary, our results show that Bro1 affects Scr1 localisation and subsequent Ght protein expression. In addition, deleting *tor1* restores the sensitivity to TOR inhibition of *bro1* mutant cells.

## Discussion

Our work positions Bro1 as an essential coordinator of chronological lifespan and TOR inhibition-mediated lifespan extension in fission yeast, *S. pombe*. Unlike cases where decreased TORC1 activity leads to lifespan extension in both normal and mutant cells through diverse mechanisms^10,20,50^, *bro1*Δ cells are short-lived and fail to modulate their survival/ageing trajectory due to TOR inhibitors such as torin1. Other identified short-lived mutants that are resistant to TOR inhibition, such as *gaf1*Δ only partially suppress the beneficial effects of torin1^17^. Our results suggest that intact endomembrane trafficking is critical for the adaptive, lifespan-extending effects of TOR suppression. This is a recuring theme in genome-wide screens we have performed with a variety of TOR inhibitors including the third-generation inhibitor, rapalink-1^37^. It will be interesting to further explore whether the different ESCRT mutants identified through our screens together with *bro1* have the same impact in ‘dampening’ the lifespan-extension effects of torin1, of other TOR inhibitors or even the lifespan extension effects through glucose^51^ or nitrogen starvation^52^.

Using Synthetic Genetic Arrays^19,38^, we identified *bro1*Δ interactions with genes involved in vesicle trafficking, vacuole fusion, and plasma membrane transporters, each central to nutrient sensing and TOR signalling. These findings agree with reports that ESCRT and HOPS/CORVET dysfunction in *S. cerevisiae* leads to vacuole fragmentation, amino acid starvation, and sensitivity to rapamycin^53^. Indeed, Bro1’s mammalian homolog ALIX binds ESCRT-III and ubiquitin, facilitating cargo sorting into multivesicular bodies (MVBs)^54,55^ and we demonstrate that *bro1*Δ cells exhibit vacuolar fragmentation, reduced amino acid reserves, implying conserved regulatory nexus across species.

Our data showcase that TOR inhibition can partially restore vacuole morphology and intracellular amino acid pools in *bro1*Δ cells. Yet, it cannot prolong lifespan, suggesting that structural rescue is insufficient for lifespan rescue without Bro1-mediated metabolic and possibly proteostatic reprogramming. This is consistent with previous observations, which highlight that TOR (particularly, TORC1) localizes on endosomes and vacuolar membranes and regulates degradation pathways^56–58^; loss of Bro1 disrupts this synergy. Interestingly, forced association of Bro1 with Psr1 (a cell membrane protein phosphatase, linked to TORC1 inhibition) in budding yeast, causes a growth defect^59^. Our proteomic analyses further reveal that Bro1 plays a central role in coordinating TOR-dependent proteome remodelling, thereby coupling vacuolar trafficking to metabolic adaptation. In wild-type cells, TOR inhibition elicited the expected suppression of ribosome biogenesis and a concomitant induction of catabolic, metabolic, and vacuolar pathways, reflecting a shift from anabolic growth toward cellular maintenance and recycling. Loss of Bro1 did not abolish ribosomal repression per se, but altered the broader proteomic response, particularly the enrichment of metabolic and vacuolar processes. This finding is consistent with our earlier observations that *bro1*Δ cells exhibit defects in vacuolar dynamics and stress resilience, despite retaining partial TOR responsiveness. Notably, *bro1*Δ cells also displayed constitutive enrichment of lipid and small-molecule metabolic pathways and elevated abundance of nutrient transporters under basal conditions, suggesting that Bro1 constrains inappropriate nutrient uptake and metabolic activity when TOR signalling is compromised. Crucially, we identify a novel functional circuit: Bro1 controls nutrient transporter homeostasis via regulation of Scr1, Ghts and Isp7. In *bro1*Δ cells, Ghts are overexpressed while Ght5 can be retained on the plasma membrane even under TOR-inhibited conditions. Our results strongly indicate that these phenomena may underpin the growth-permissive, TOR-resistant phenotype of the *bro1* mutant cells. We further link Bro1 to Scr1 ubiquitination and mislocalization, suggesting that Bro1 supports a novel nucleus-to-vacuole signalling route for transcriptional regulation. This aligns with Bro1’s recruitment of ubiquitin hydrolases like Doa4 in *S. cerevisiae*, ensuring controlled deubiquitination of cargoes destined for MVBs^34,60,61^.

Our RNA-seq analyses reveal that *bro1*Δ cells mount a more pronounced transcriptional response under TORC1-specific inhibition (caffeine-rapamycin) compared to *wt*, including downregulation of ribosome biogenesis and 5.8S rRNA processing at late timepoints, pointing to maladaptive proteostasis shifts. This aligns with previous studies, demonstrating that the endolysosomal membrane trafficking system can drive TORC1 towards anabolism and growth^62^ and that the late endosome is essential for TORC1 function^63^. Scr1 and hexose transporter deregulation and vacuolar fragmentation occur in *bro1*Δ cells together. This suggests a multi-tiered coordination failure in response to nutrient stress. From a cell biological perspective, the altered enrichment of vacuolar transport, vesicle-mediated trafficking, and autophagy-associated proteins in *bro1*Δ cells reinforces a model in which Bro1 coordinates ESCRT-driven endosomal sorting and vacuolar dynamics with proteome turnover. Defective execution of these processes would be expected to compromise both protein quality control and metabolite recycling, processes that are increasingly recognised as central determinants of cellular ageing. Consistent with this view, the constitutive upregulation of lipid metabolic pathways and nutrient transporters observed in *bro1*Δ cells suggests a failure to appropriately restrain anabolic and uptake programmes, even under conditions that normally favour maintenance and longevity. This is further supported by studies indicating that deregulated transporter recycling and excess nutrient uptake impair vacuolar-autophagic signalling and lifespan^64–66^.

Together, our findings place Bro1 at the intersection of endosomal sorting, nutrient transporter regulation, transcriptional modulation via Scr1, and TOR signalling output, a framework with potential relevance to diverse higher eukaryotic systems where ALIX-family proteins regulate receptor trafficking and signal transduction^67,68^. Future work should delineate direct physical interactions between Bro1, Scr1, E3 ligases, AMPK/Ssp2, and TORC1 components, while exploring whether mitochondrial ROS and autophagic flux are impacted by Bro1 status, given their established roles in lifespan modulation^69^. Our data also align with emerging views in cancer biology that lysosomal and endosomal functions can underlie resistance to mTOR inhibitors^70,71^, suggesting targeting Bro1-mediated trafficking may enhance therapeutic efficacy.

In summary, Bro1 is a central integrator of nutrient trafficking, transporter expression, vacuolar integrity, TOR signalling, and lifespan. Its absence decouples TOR inhibitor response from lifespan benefits, providing mechanistic insight into the cellular coordination necessary for longevity and highlighting conserved pathways for therapeutic targeting.

## Materials and Methods

### Strains and media

Strains used in the study are presented in Table S7. The wild-type strain is 972^-^. Cells were grown on YES and EMM2 media as stated in the main text and legends.

### Chronological lifespan

Cells were grown in EMM2 media as previously described ^10^. When cultures reached a stable maximal density, cells were harvested, serially diluted, and incubated on YES plates. The measurement of colony-forming units (CFUs) was taken at the beginning of the lifespan curve (time point 0: 100% cell survival). Lifespans are calculated from three independent cultures, with each culture measured three times at each time point. To determine the chronological lifespan when TOR is inhibited, 5 μM Torin1 was added to rapidly proliferating cell cultures at OD_600_ 0.5 which were then grown to stationary phase, and lifespan was recorded as described above. AUCs were measured with ImageJ ^72^ for all experimental repeats using lifespans curves on the linear scale for percentages of survival. Log-rank testing is used for p-value generation for the obtained survival curves.

### Quantitative Amino Acid Profiling

Metabolite extraction and amino acid analysis was performed using hydrophilic interaction chromatography-tandem mass spectrometry (HILIC-MS/MS) as described in^45,48,73^. Cell numbers were determined with a Beckman Z-series coulter counter to ensure equal cell amounts for extractions. Identification of 19 proteogenic amino acids was obtained by comparison of retention time and fragmentation patterns with commercial standards. To quantify the exact amino acids, we included cell-free, ^15^N, ^13^C-labelled amino acids (CNLM-6696-PK, Cambridge Isotope Laboratories) as an internal standard. Additionally, a calibration curve was generated using serial dilutions of unlabeled standards. Amino acid measurements were performed at a flow rate of 0.6 ml/min for 12 minutes. The analysis was undertaken using an Agilent Infinity 1290 LC system with ACQUITY UPLC BEH amide columns (Waters Corporation, Manchester, United Kingdom) (pore diameter 130 Å, particle size 1.7 μm, internal diameter 2.1, column length 100 mm) coupled to an Agilent 647 Series Triple Quadrupole LC/MS mass spectrometer operating in selected reaction monitoring mode. Cysteine was excluded from analysis due to its low stability.

#### Proteomes

Samples were mechanically lysed in 200 µl of 8 M urea, 0.1 M ammonium bicarbonate (Geno/Grinder, SPEX). Semi-automated sample preparation was performed in 96-well format, using in advance prepared stock solution plates stored at -80°C as previously described^74^. To reduce disulfide bonds, 5 mM dithiothreitol (final concentration) was added to each sample, followed by incubation at 30°C for 1 hr. Next, 20 µL of 0.1 M iodoacetamide were added to each sample to acetylate previously reduced thiol groups. After an incubation of 30 min at 20 °C in the dark, samples were diluted by adding 450 µL of 0.1 M ABC + Benzonase. After an incubation at 37°C for 30 min, 500 µl of sample were transferred to prepared 2 ug/well trypsin/lysC plates. Samples were incubated overnight at 37°C. Digestion was stopped by the addition of 20% formic acid (FA). Digestion mixture was cleaned-up by using MCX 2 mg Sorbens plates (Oasis MCX 96-well). The eluted peptides were diluted by adding 75 µl 0.1% formic acid. Peptides were analyzed on a Bruker timsTOF HT mass spectrometer coupled to an Infinity II LC system (Agilent). 5 µg of the sample were chromatographically separated on a Phenomenex Luna®Omega column (1.6 μm C18 100Å, 30 × 2.1 mm) heated to 50°C, using a flow rate of 0.5 ml/min where mobile phase A & B were 0.1% formic acid in water and 0.1% formic acid in acetonitrile, respectively. The LC gradient ran as follows: 3% to 36% B in 5 min, increase to 80% B at 0.8 mL over 0.5 min, which was maintained for 0.2 min and followed by equilibration with starting conditions for 2 min. For diaPASEF MS acquisition, the electrospray source (Bruker VIP-HESI, Bruker Daltonics) was operated at 3000 V of capillary voltage, with a nebulizer pressure of 4.5 bar, 10.0 l/min of drying gas, and 240°C drying temperature. Sheath gas temperature and flow were 450°C and 4.8 l/min respectively. The diaPASEF windows scheme was as follows: we sampled an ion mobility range from 1/K0 = 1.30 to 0.7 Vs/cm2 using ion accumulation time and ramp time of 72 ms in the dual TIMS device and a mass range of 400 to 1125 Da. Each cycle estimates a time of 0.7 s. The collision energy was lowered as a function of increasing ion mobility from 59 eV at 1/K0 = 1.6 Vs/cm2 to 20 eV at 1/K0 = 0.6 Vs/cm2. TIMS device was linearly calibrated using three Agilent ESI-L Tuning Mix ions (m/z, 1/K0: 622.0290, 0.9915 Vs/cm²; 922.0098, 1.1986 Vs/cm²; and 1221.9906, 1.3934 Vs/cm²). The mass spectrometry data was analysed with DIA-NN (version 1.8.1)^75^. In brief, an in-silico spectral library was generated from Uniprot reference proteome UP000002485 with a gene count of 5206 (downloaded 2025-07-04). Collected spectra were annotated and signals quantified using default DIA-NN settings, except for mass accuracy which was set at 15 for both MS1 and MS2 and a scan window size of 7.

#### RNA-seq

Wild-type (*972 h-*) and *bro1* mutant cells were grown on YES media in the presence or absence of Torin1 (2 µM) or combination of caffeine/rapamycin (10mM caffeine and 100µM rapamycin) at timepoints as indicated in main text and legends. Cells were then harvested at early exponential growth phase (OD_600_ 0.5), and total RNA was isolated by hot-phenol extraction^76^. RNA quality was assessed on a Bioanalyzer instrument (Agilent), treated with DNase (Turbo DNA-free, Ambion) and subsequently RNA was treated with a beta version of Ribo-Zero Magnetic Gold Kit Yeast (Epicentre) to deplete rRNAs. RNA-seq libraries were prepared from rRNA-free RNA using a strand-specific library preparation protocol and sequenced on an Illumina HiSeq instrument. Standard sequence data analysis was carried out as previously described ^37,77^, using 150-bp paired reads (GEO accession number: GSE295379).

#### Microscopy, vacuolar staining and measurements

To determine cell size, control and drug-treated cells were fixed in 4% formaldehyde for 10 min at room temperature and stained with Calcofluor (50 mg/ml). Microscopy was performed using a DAPI filter for Calcofluor detection on an EVOS5000 system using 40x or 60x lenses. At least 100 septated cells were counted and analyzed for each condition with results analyzed in R. GFP microscopy has been conducted as described in ^37^.Vacuolar labeling was performed as described in ^78^. For vacuole visualization and size quantification, cells were harvested at mid-log phase and resuspended in 10 mM HEPES buffer (pH 7.4) containing 5% glucose. Cells were incubated at 30°C for 15–30 min with CMAC (7-amino-4-chloromethylcoumarin; Invitrogen) to label the vacuole lumen, washed once with 1× PBS and immediately imaged using a 63× oil-immersion objective on a Leica SPE or SPE2 confocal microscope. Images were acquired in a single focal plane using LAS AF software (Leica Microsystems). Vacuole lumen areas were automatically measured for at least 1000 cells per condition using ImageJ. Vacuole diameter was calculated and the mean vacuole diameter was used as the vacuole size for each population.

#### Western blotting

For protein preparations, cells were diluted in RIPA buffer supplemented with protease and phosphatase inhibitors (Sigma cocktails 1 and 2), together with glass beads. Cells were lysed in a Fastprep-24 machine (MP Biomedicals). Anti-GFP antibody (Roche #11814460001) have been used for GFP detection (1/2000 dilutions). Anti-Ubiquitin antibody (Proteintech #80992-1-RR) has been used for Ubiquitinated Lysine detection (1/120000). For detection in Fig.5B we used anti-rabbit IRDye 680LT conjugated secondary antibody ((1/10000 dilution, #925-68023, Licor). For all other detections, we used the anti-mouse HRP-conjugated antibody (1/5000 dilutions) with the Supersignal West Pico PLUS detection system (thermo scientific, #34580) according to the manufacturer’s protocol.

#### Immunoprecipitations

Immunoprecipitation has been performed using DiaMag protein A-coated magnetic beads (Diagenode ChIP kit #C01010055). 40ug protein extracts with 0.2 µg anti-GFP antibody in a total volume of 20 µL RIPA buffer containing protease and phosphatase inhibitors (RIPA+PPI) were incubated overnight at 4°C on a spinning wheel. 30 µL of beads were washed with RIPA+PPI and left in ice for 5 min. The protein-antibody mix from the overnight incubation was added to the beads and incubated on spinning wheel at 4°C for 2hrs. Supernatant was removed using MagRack and collected (“Flow through”). Beads were washed twice with 100 µL RIPA+PPI. 25 µL sample buffer was added and samples were boiled at 80°C for 10 min. Samples were processed for western blotting.

## Acknowledgements and Funding

This work was supported by funding to C.R. from the Biotechnology and Biological Sciences Research Council [Research grant number: BB/V006916/1 and BB/V006916/2] and the Medical Research Council [Grant number: MR/W001462/1 and MR/W001462/2). D.S was supported by the Francis Crick Institute which receives its core funding from Cancer Research UK (CC001051), the UK Medical Research Council (CC001051), and Wellcome (CC001051). We acknowledge the German Research Foundation (DFG, Deutsche Forschungsgemeinschaft) – SFB/TRR 186 (Project number 278001972), to MR and supporting N.B. as well as the Ministry of Education and Research (BMBF), as part of the National Research Initiative ‘Mass Spectrometry in Systems Medicine’ (MSCoreSys), under grant agreement number 01EP2201 (to M.R.) and 16LW0239K (to M.M.). This work was supported by awards to S.E.M (ORCID:0000-0003-4385-4957) from the European Union’s Horizon 2020 research and innovation programme (BATCure, grant No 666918), and the USA Children’s Brain Disease Foundation. Microscopy was performed at the Light Microscopy Core Facility, UCL GOS Institute of Child Health supported by the NIHR GOSH BRC award 17DD08. The facility is funded in part by an award from the NIHR Great Ormond St Hospital Biomedical Research Centre. All research at UCL Great Ormond Street Institute of Child Health is made possible by the NIHR Great Ormond Street Hospital Biomedical Research Centre. The views expressed are those of the authors and not necessarily those of the NHS, the NIHR or the Department of Health. We are grateful to Shigeaki Saitoh for providing strains used in this study.

## Authors’ contributions

Conceptualization, C.R.; Methodology, C.R. Investigation, J.K., K.N., A.D., J.C-R., R.I., O. X., J-P.A., D.S., S.E.M., N.B., M.R., M.M. and C.R.; Formal Analysis, C.R., J.K., K.N., writing, C.R., J.K., A.D.; Funding Acquisition, C.R., S.E.M., M.R., M.M., P.T.; Supervision, C.R.

## Supplemental Figures

**Figure S1.**
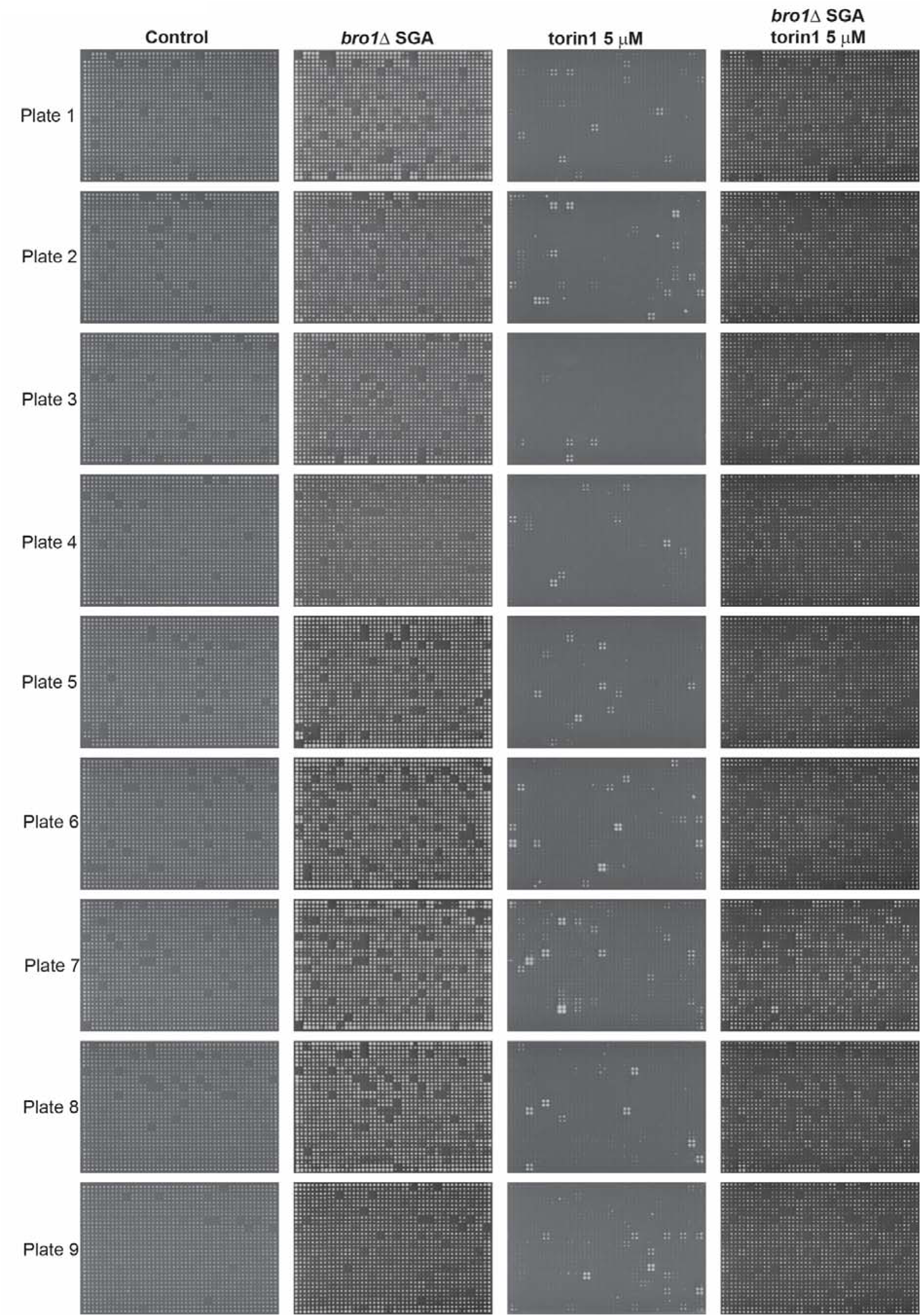
Gene-gene, gene-drug and gene-gene-drug raw image data acquisition. Representative images of plates from *bro1*Δ Synthetic Genetic Arrays, torin1 resistance screens and combinations of the two. Note the few resistant strains recovered from the torin1 screen and the effect of *bro1*Δ in the resistance phenotype of the fission yeast deletion library.

**Figure S2.**
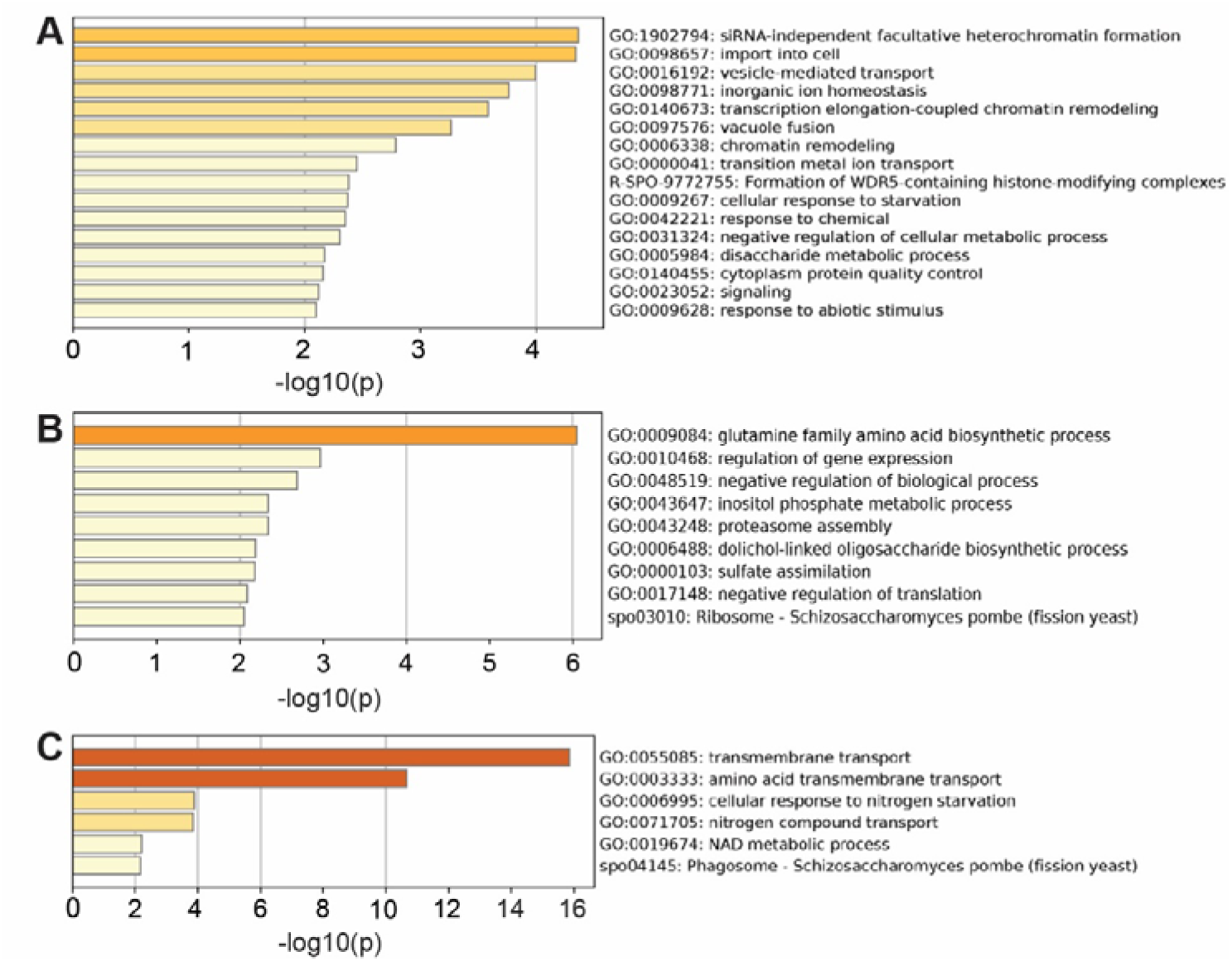
**A.** Gene ontology (GO) enrichments for *bro1* negative interactions. **B.** GO enrichments for *bro1* positive interactions. **C.** GO enrichments for predicted Bro1 protein-protein interactions using the Pint tool.

**Figure S3.**
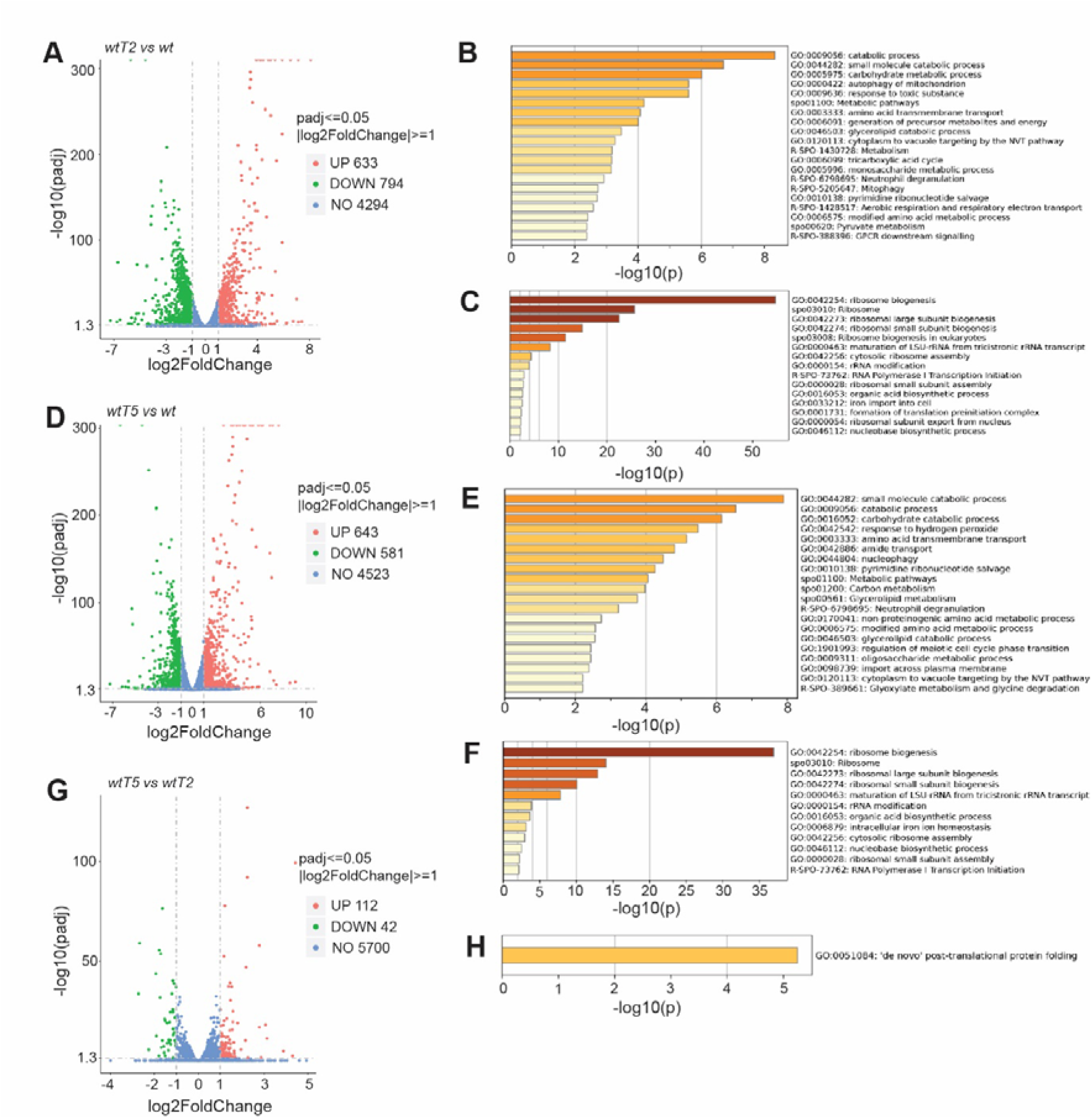
Gene expression analyses for torin1 treatments and untreated conditions for *wt* cells. **A.** Volcano plot for 2-fold differentially expressed genes between torin1 2 hr-treated and untreated *wt* cells. **B.** GO enrichments for upregulated genes in A. **C.** GO enrichments for downregulated genes in A. **D.** Volcano plot for 2-fold differentially expressed genes between torin1 5 hr-treated and untreated *wt* cells. **E.** GO enrichments for upregulated genes in D. **F.** GO enrichments for downregulated genes in D. **G.** Volcano plot for 2-fold differentially expressed genes between torin1 5 hr-treated and torin1 2 hr-treated *wt* cells. **H**. GO enrichments for upregulated genes in G. No significant enrichment is observed for the 42 downregulated genes.

**Figure S4.**
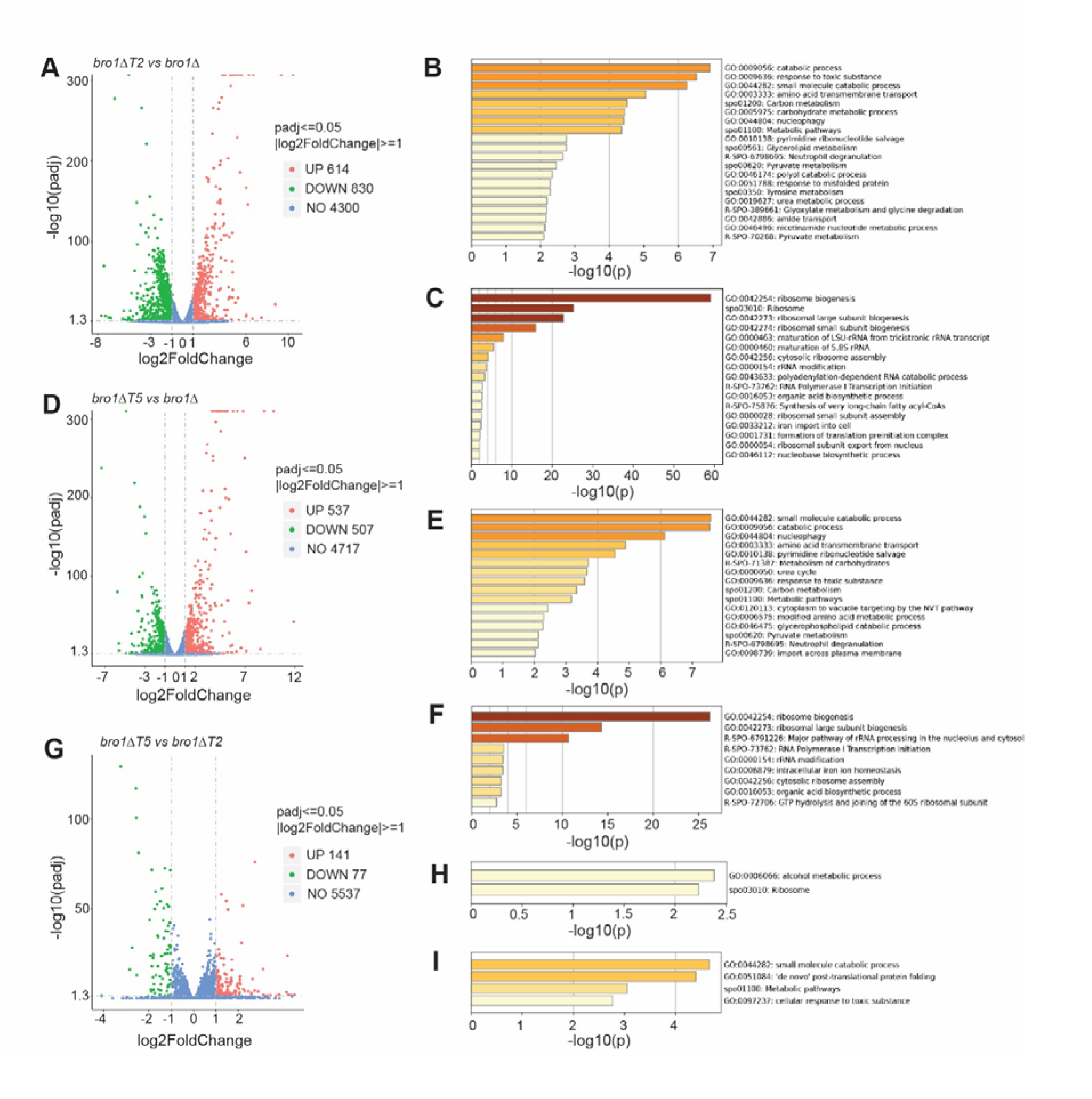
Gene expression analyses for torin1 treatments and untreated conditions for *bro1*Δ cells. **A.** Volcano plot for 2-fold differentially expressed genes between torin1 2 hr-treated and untreated *bro1*Δ cells. **B.** GO enrichments for upregulated genes in A. **C.** GO enrichments for downregulated genes in A. **D.** Volcano plot for 2-fold differentially expressed genes between torin1 5 hr-treated and untreated *bro1*Δ cells. **E.** GO enrichments for upregulated genes in D. **F.** GO enrichments for downregulated genes in D. **G.** Volcano plot for 2-fold differentially expressed genes between torin1 5 hr-treated and torin1 2 hr-treated *bro1*Δ cells. **H**. GO enrichments for upregulated genes in G. **I.** GO enrichments for downregulated genes in G.

**Figure S5.**
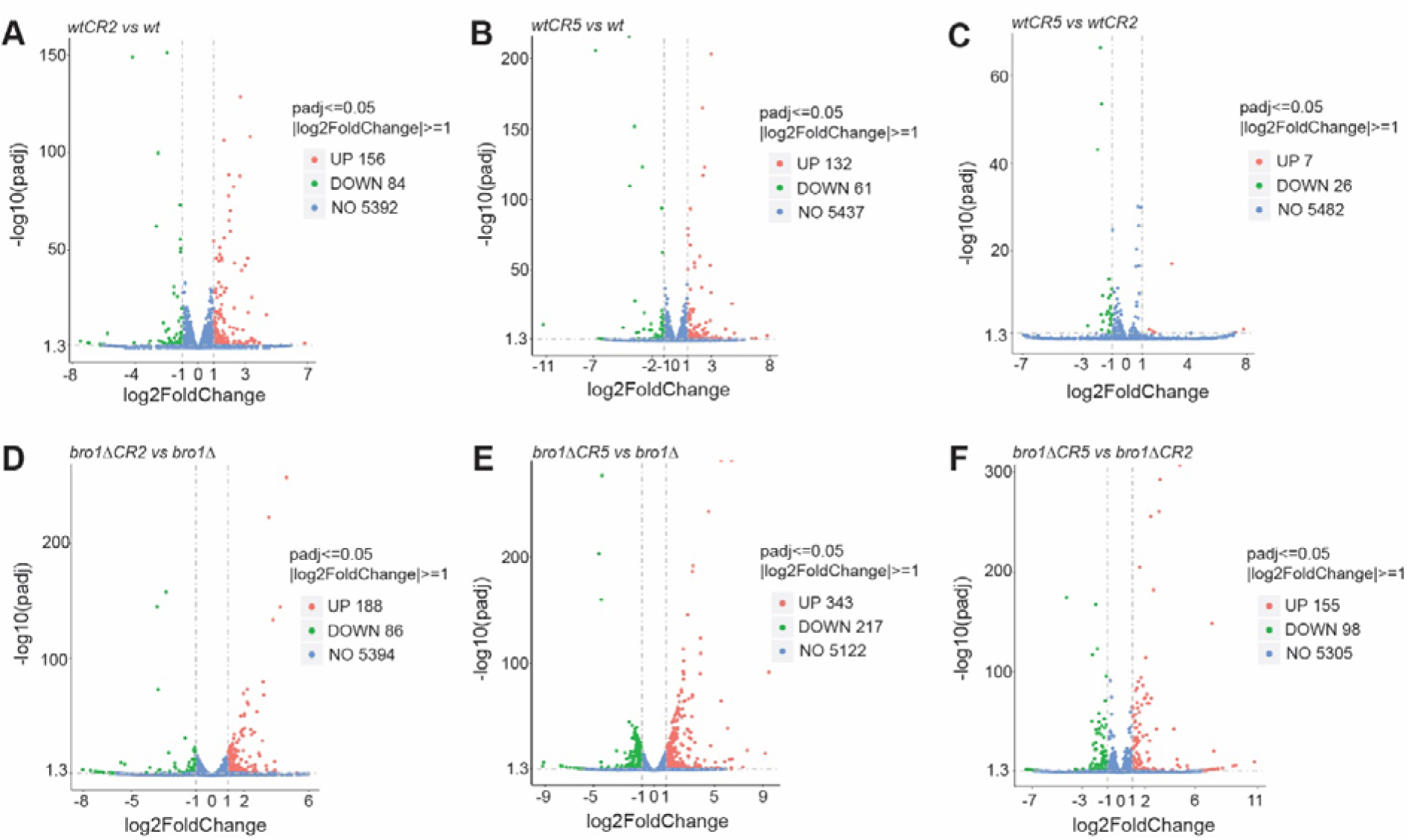
Volcano plots showcasing the relatively small number of genes that are differentially expressed at a 2-fold cutoff for *wt* cells (**A-C**) and *bro1*Δ (**D-F**) cells treated with caffeine and rapamycin (CR) for 2 hrs (CR2) or 5 hrs (CR5). Using this cutoff there are significant GO enrichments (see main text for more information).

**Figure S6.**
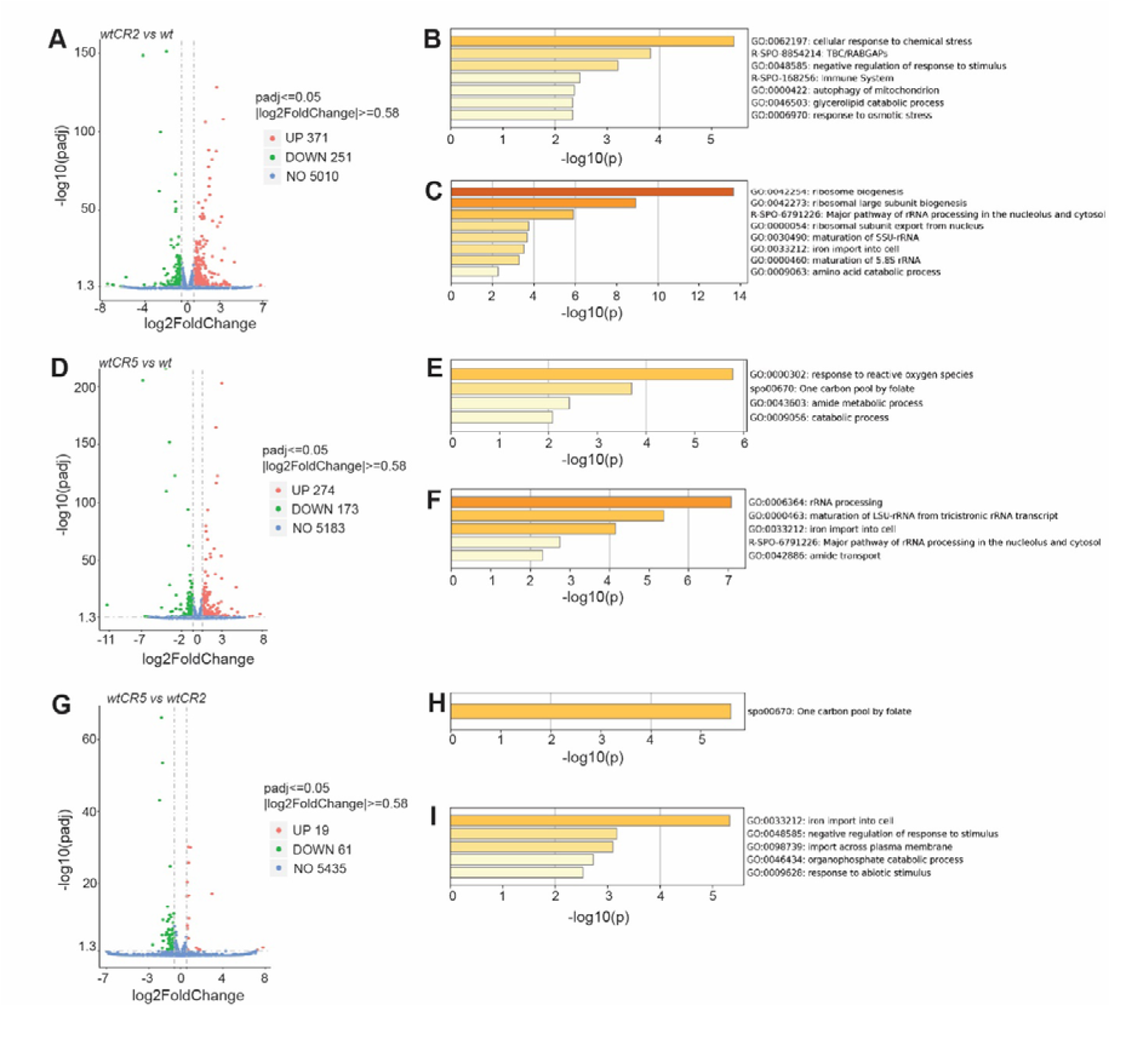
Gene expression analyses for caffeine-rapamycin treatments and untreated conditions for *wt* cells. **A.** Volcano plot for 1.5-fold differentially expressed genes between caffeine-rapamycin 2 hr-treated and untreated *wt* cells. **B.** GO enrichments for upregulated genes in A. **C.** GO enrichments for downregulated genes in A. **D.** Volcano plot for 1.5-fold differentially expressed genes between caffeine-rapamycin 5 hr-treated and untreated *wt* cells. **E.** GO enrichments for upregulated genes in D. **F.** GO enrichments for downregulated genes in D. **G.** Volcano plot for 1.5-fold differentially expressed genes between caffeine-rapamycin 5 hr-treated and caffeine-rapamycin 2 hr-treated *wt* cells. **H**. GO enrichments for upregulated genes in G. **I.** GO enrichments for downregulated genes in G.

**Figure S7.**
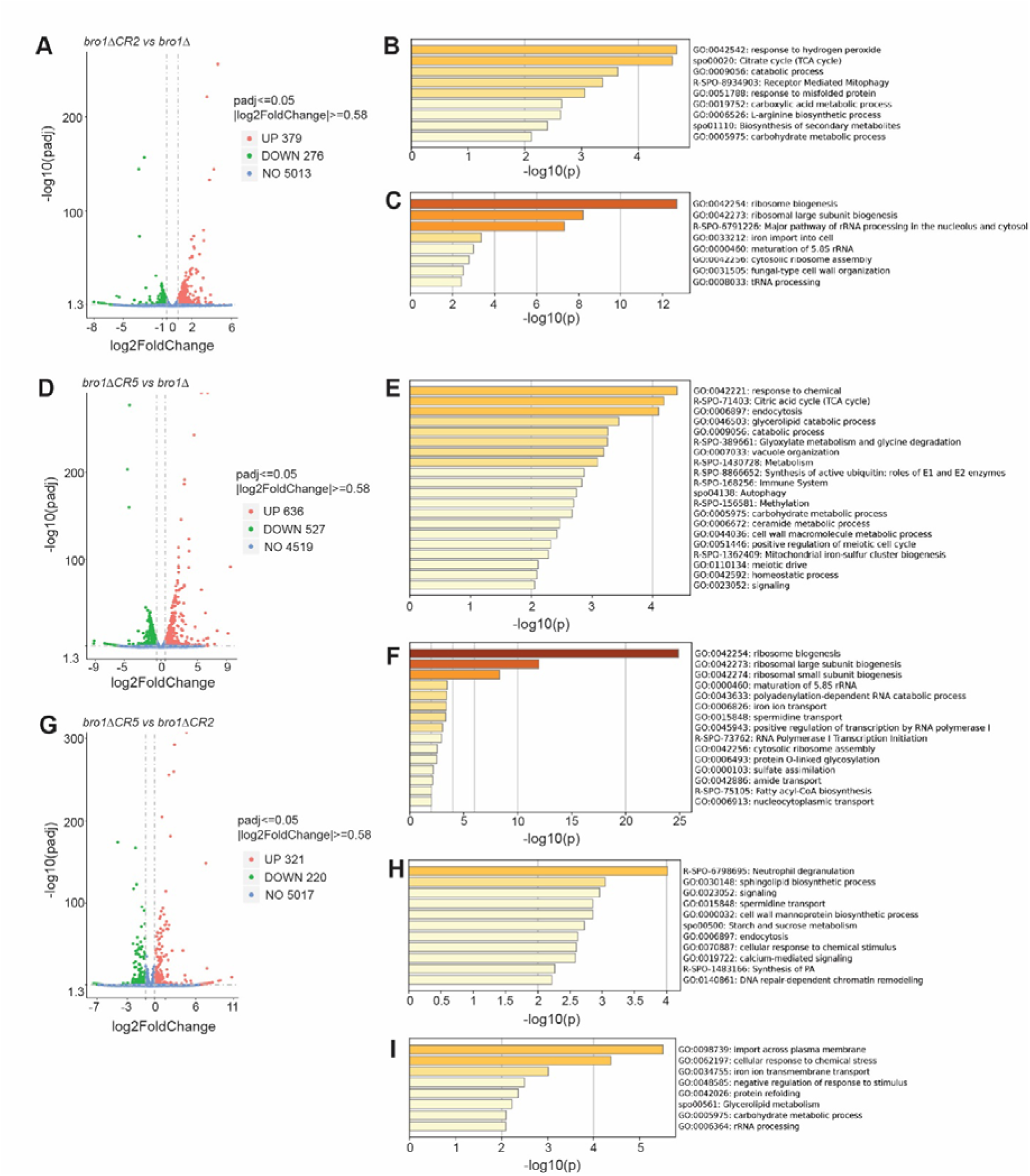
Gene expression analyses for caffeine-rapamycin treatments and untreated conditions for *bro1*Δ cells. **A.** Volcano plot for 1.5-fold differentially expressed genes between caffeine-rapamycin 2 hr-treated and untreated *bro1Δ* cells. **B.** GO enrichments for upregulated genes in A. **C.** GO enrichments for downregulated genes in A. **D.** Volcano plot for 1.5-fold differentially expressed genes between caffeine-rapamycin 5 hr-treated and untreated *bro1Δ* cells. **E.** GO enrichments for upregulated genes in D. **F.** GO enrichments for downregulated genes in D. **G.** Volcano plot for 1.5-fold differentially expressed genes between caffeine-rapamycin 5 hr-treated and caffeine-rapamycin 2 hr-treated *bro1Δ* cells. **H**. GO enrichments for upregulated genes in G. **I.** GO enrichments for downregulated genes in G.

**Figure S8.**
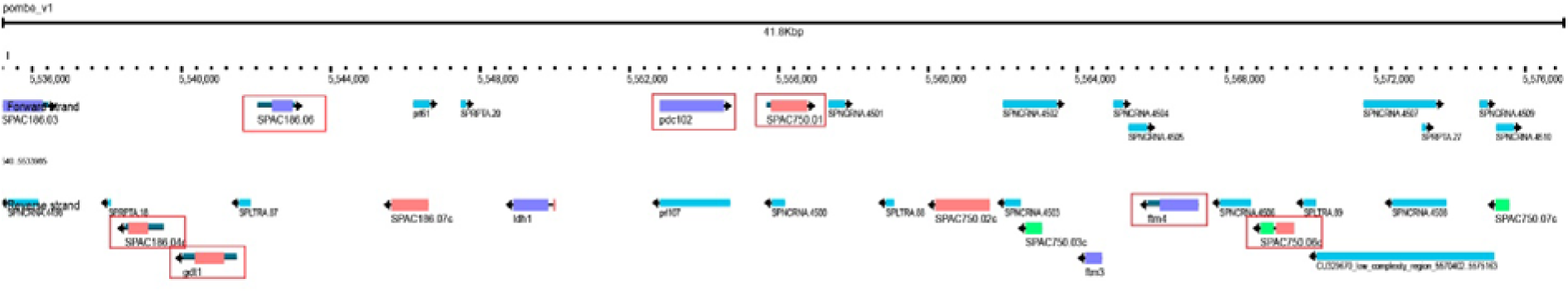
Genome browser capture of fission yeast chromosome I (area 5,536,000-5,576,000). Genes in boxes are upregulated at least 1.5-fold in *bro1*Δ cells compared to *wt*.

## Supplemental Tables

**Table S1**

Genetic interaction values for 2,993 deletion mutants with *bro1* obtained through Synthetic Genetic Arrays (SGAs) as previously described^19,38^ (see main text for more information).

**Table S2**

Genetic interaction values for 3,101 deletion mutants with *bro1* in the presence of torin1 obtained through Synthetic Genetic Arrays (SGAs).

**Table S3**

Top 100 hits of predicted physical interactions of Bro1 using the tool Pint^43^

**Table S4**

List of 2-fold differentially expressed genes between *bro1*Δ and *wt* cells.

**Table S5**

Gene expression fold-change values between *bro1*Δ and *wt* cells accompanied with p-values for all genes obtained. All RNA-seq data are deposited and available on Gene Expression Omnibus (GEO accession number: GSE295379).

**Table S6**

Summary of proteomics results obtained from this study.

**Table S7**

List of fission yeast strains used in this study.

